# Nuclear HMGB1 protects from non-alcoholic fatty liver diseases through negative regulation of liver X receptor

**DOI:** 10.1101/2021.02.17.431446

**Authors:** Jean Personnaz, Enzo Piccolo, Alizée Dortignac, Jason S. Iacovoni, Jérôme Mariette, Arnaud Polizzi, Aurélie Batut, Simon Deleruyelle, Romain Paccoud, Elsa Moreau, Frédéric Martins, Thomas Clouaire, Fadila Benhamed, Alexandra Montagner, Walter A. Wahli, Robert F. Schwabe, Armelle Yart, Isabelle Castan-Laurell, Catherine Postic, Cédric Moro, Gaelle Legube, Chih-Hao Lee, Hervé Guillou, Philippe Valet, Cédric Dray, Jean-Philippe Pradère

## Abstract

Dysregulations of lipid metabolism in the liver may trigger steatosis progression leading to potentially severe clinical consequences such as non-alcoholic fatty liver diseases (NAFLD). Molecular mechanisms underlying liver lipogenesis are very complex and fine-tuned by chromatin dynamics and the activity of multiple key transcription factors. Here, we demonstrate that the nuclear factor HMGB1 acts as a strong repressor of liver lipogenesis during metabolic stress in NAFLD. Mice with liver-specific *Hmgb1*-deficiency display exacerbated liver steatosis and hepatic insulin resistance when subjected to a high-fat diet or after fasting/refeeding. Global transcriptome and functional analysis revealed that the deletion of *Hmgb1* gene enhances LXRα activity resulting in increased lipogenesis. HMGB1 repression is not mediated through nucleosome landscape re-organization but rather via a preferential DNA occupation in region carrying genes regulated by LXRα. Together these findings suggest that hepatocellular HMGB1 protects from liver steatosis development. HMGB1 may constitute a new attractive option to therapeutically target LXRα axis during NAFLD.

## Introduction

Along the epidemic of obesity, non-alcoholic fatty liver disease (NAFLD) is progressing worldwide, affecting nearly 25% of the world-wide adult population (*1*) and generating numerous complications such as liver insulin resistance, non-alcoholic steatohepatitis and hepatocellular carcinoma (*2*). Liver steatosis consists in ectopic lipid storage within the hepatocytes, which aims at buffering circulating lipids and thus preventing lipotoxicity in different organs. Mechanisms underlying lipogenesis (from lipid uptake to lipid esterification and *de novo* lipogenesis) are extremely complex and consist in a subtle orchestration of the actions of different transcription factors (TFs) in close coordination with chromatin dynamics (*3*).

Among TFs involved in liver lipogenesis regulation, Liver X Receptors (LXRs) are members of the nuclear hormone receptor superfamily and are among the most central/dominant actors in this process. LXRs consist in two isotypes that share a very high homology but differ in their tissue expression profile. While LXRα (NR1H3) is mainly expressed in metabolic tissues (liver, adipose tissues), LXRβ (NR1H2) is expressed ubiquitously (*4*). In the context of dyslipidemia or fasting/refeeding conditions and after activation by certain lipid species (*5*), LXRs directly coordinate, in a duo with its obligate partner, retinoic acid receptor (RXR), the expression of numerous key enzymes involved in cholesterol and lipid metabolism (*Abcg5*, *Abcg8*, *Fasn*, *Scd-1*), but are also capable to modulate indirectly the lipogenesis through the regulation of other key TFs like SREBP1c, ChREBP or PPARγ (*4*, *6*, *7*) that are also involved in the lipogenic transcription program. The current consensus on liver lipogenesis is that there is a hierarchical interplay between all TFs involved, where LXR is a very central piece; SREBP1 and ChREBP are crucial downstream key players while PPARγ’s role appears more supportive (*8*). LXRs activity is subtly regulated by the interaction with the nuclear receptor co-repressors (NCoR) or the nuclear receptor coactivators protein complex (*8*) upon specific agonist activation. Recent evidences are now showing the emerging role of some methylase/demethylase enzymes in the modulation of LXR activity through the chromatin packaging and subsequent availability, adding one more complex layer of regulation (*9*, *10*).

Global knockout of LXRs induces a severe reduction of liver lipid synthesis in wild type mice and could even prevent liver steatosis in ob/ob mice (*11*–*13*). LXRα deletion knockout leads to a drastic down-regulation of *Srebf1* expression associated with a reduced lipogenesis (*6*). Moreover LXRs agonist treatment increases plasma and hepatic TG in mice and humans (*14*, *15*) supporting a key role of LXRs in fatty acid synthesis and liver steatosis progression. Therapeutic targeting of LXRs is still challenging as adverse effects have been described (*15*) and more insights regarding LXRs upstream regulators may be helpful to design novel therapeutic avenues.

HMGB1 belongs to the family of high mobility group proteins, which after the histones represents the most abundant proteins in the nucleus. In recent years, HMGB1 has also been scrutinized for its role in the extracellular cellular compartment as a potent inflammatory factor, notably during sterile inflammation (*9*). Originally however, HMGB1 has been known for its role in the nucleus (*17*) as a protein capable of binding chromatin on unspecific domains (*18*) in a very dynamic manner (*19*). HMGB1 may affect several biological functions such as VDJ recombination, DNA repair (*20*), chromatin assembly and gene transcription through different mechanisms, such as DNA bending/looping, nucleosome formation (*21*, *22*), interaction with the transcription machinery including TFs themselves (*19*, *23*–*25*). A very recent report depicts nuclear HMGB1 as an even more versatile factor able to bind to topologically-associated domains or RNA directly to regulate proliferation or senescence programs (*26*). In cultured cells, while HMGB1 deletion leads to minor changes in histone numbers, it results in notable changes of the RNA pool (*22*), in local chromatin remodeling (*27*) or the global transcriptome (*26*). However only a sparse number of studies have been carried out *in vivo* (*27*). The global ablation of *Hmgb1* generates a severe phenotype with perinatal mortality (*28*), likely due to a defective glucocorticoid signaling leading to a poor utilization of hepatic glycogen and resulting in a lethal hypoglycemia, whereas hepatocyte-specific HMGB1 ablation did not have a major impact under homeostatic conditions (*29*). Thus, in this context it seems particularly relevant to explore the role of nuclear HMGB1 *in vivo* especially during metabolic stress, where the dynamics of the chromatin are critical to orchestrate the activity of key TFs and gene transcription programs in order to buffer stress mediators and maintain whole-body homeostasis.

Here, we unveiled the important role of HMGB1 in the repressive effect of LXRs, in particular LXRα, during metabolic stress, as demonstrated by increased liver steatosis and an alteration of the hepatic insulin in hepatocyte-specific *Hmgb1* knockout (HMGB1^ΔHep^) mice subjected to either a high-fat diet (HFD) or a fasting-refeeding (F/R) challenge. *In vitro* assays further confirmed the repressive action that HMGB1 exerts on LXRα activity. Taken together, our data reveal a novel role of HMGB1 in alleviating liver steatosis through the repression of LXRα during metabolic stress.

## Results

### Hepatic deletion of Hmgb1 increases liver steatosis during metabolic stress

*Hmgb1* hepatocyte-specific knockout mice (HMGB1^ΔHep^) under chow diet (CD) feeding display no major changes in liver transcriptome and no drastic phenotype of glycogen utilization compared to control mice (HMGB1^fl/fl^) (*29*), contrasting findings from the global *Hmgb1* knockout on metabolism, possibly due to particular functions during development (*28*). This prompted us to clarify the precise function of HMGB1 in liver metabolism by studying the role of HMGB1 as a potential regulator of global and/or hepatic energy metabolism in adult mice using a careful characterization of HMGB1^fl/fl^ and HMGB1^ΔHep^ mice subjected to metabolic stress. A complete metabolic checkup in adult mice upon CD, showed that deletion of *Hmgb1* in hepatocytes (fig. S1A) did not affect circulating levels of HMGB1 (fig. S1B), serum liver enzyme levels (fig. S1C), body weight (fig. S1D), lean/fat mass ratio (fig. S1E) fasting blood glucose levels and glucose homeostasis (fig. S1F) nor generated any changes in hepatic lipid contents (fig. S1G) or in food consumption and other parameters assessed by indirect calorimetry (not shown). However, a high-throughput real-time qPCR gene expression profiling targeting metabolic pathways revealed that many key genes involved in lipid metabolism and lipogenesis, such as *Cd36*, *Fasn* or *Acly*, were upregulated in the liver of HMGB1^ΔHep^ mice compared to HMGB1^fl/fl^ mice (fig. S1H). Collectively, these data suggest, while supporting conclusions from a previous report (*29*) on the minor role of HMGB1 in systemic and liver metabolic homeostasis, that its function might become relevant in the setting of metabolic stress. To test this hypothesis, HMGB1^fl/fl^ and HMGB1^ΔHep^ mice were subjected to a high-fat diet feeding (HFD60%). After 12 weeks of this regimen, HMGB1^fl/fl^ control mice showed the expected weight gain and glucose metabolism deterioration compared to mice fed CD (not shown). In this context, after HFD60%, both genotypes displayed similar weight gain (fig. S2A) and similar fat mass (fig. S2B) and shared identical physiological parameters (food intake, respiratory quotient, physical activity) (fig. S2C-E). However, HMGB1^ΔHep^ mice exhibited a significant increase in Oil Red-O staining (**Fig. 1A**) and in liver lipid content, especially cholesterol ester, compared to control mice (**Fig. 1B**). In addition, mRNA expression analysis revealed a drastic upregulation of key genes involved in liver lipid metabolism and lipogenesis such as *Cd36*, *Fasn, Scd-1*, *Pnpla3, Adrp47* or *Lxrα* (**Fig. 1C**) in livers from HMGB1^ΔHep^ mice compared to control littermates. To further challenge the lipogenic pathway using a more acute nutritional setting without confounding effects related to a 12 week-HFD, HMGB1^fl/fl^ and HMGB1^ΔHep^ mice were subjected to a 6 hour-fast and an 8 hour-chow diet refeeding (F/R) experiment. Similar to HFD, hepatic lipid accumulation in HMGB1^ΔHep^ mice was notably more pronounced compared to control mice, as supported by a drastic increase of Oil Red-O staining on liver sections (**Fig. 1D**), of hepatic lipid levels (**Fig. 1E**) and lipogenic gene expression (**Fig. 1F**) in liver biopsies from HMGB1^ΔHep^ mice compared to HMGB1^fl/fl^ mice. To confirm the HMGB1^ΔHep^ mice phenotype, several other diets designed to challenge the hepatic lipogenesis were implemented, such as 24 week HFD, 8 week-choline deficient-HFD and a 12 week-high fat-high fructose diet, all showing a consistent and more severe liver steatosis in HMGB1^ΔHep^ mice compared to HMGB1^fl/fl^ mice (fig. S3A-C). These results indicate that under several steatosis-promoting regimens, *Hmgb1* deletion in hepatocytes is associated with a more active liver lipogenesis, suggesting that HMGB1 might play a repressive role on liver lipid synthesis, thereby preventing steatosis.

**Fig. 1.**
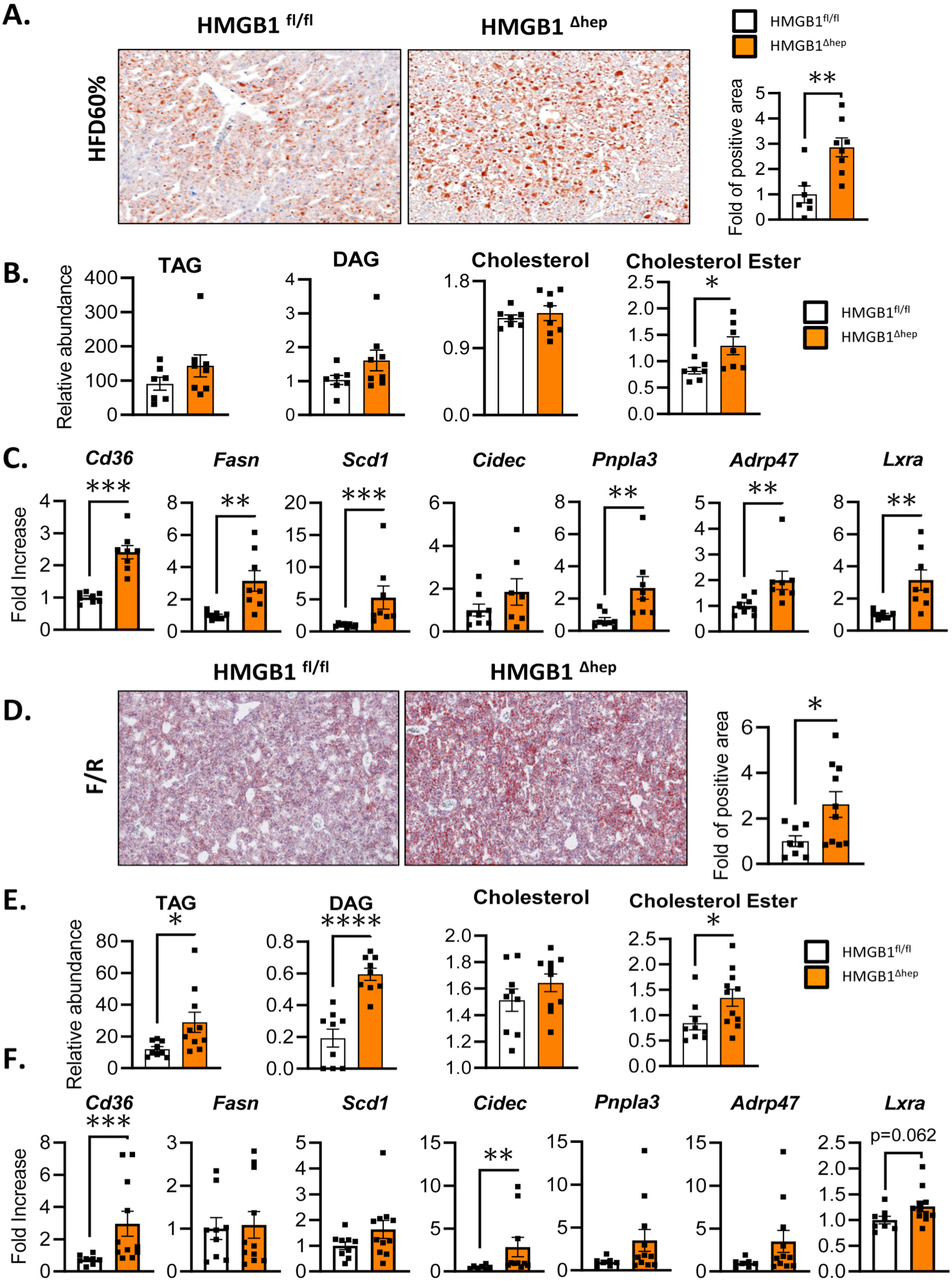
Hepatocyte specific *Hmgb1* deleted mice on HFD or after fasting/refeeding challenge exhibit a severe liver steatosis. **(A)** Oil Red-O staining on liver section with quantification, (**B**) neutral lipid analysis and (**C**) mRNA expression of hepatic steatosis markers from liver biopsies of HMGB1^fl/fl^ and HMGB1^ΔHep^ mice subjected to 12-week HFD. (**D**) Oil Red-O staining on liver section with quantification, (**E**) neutral lipid analysis and (**F**) mRNA expression of hepatic steatosis markers from liver biopsies of HMGB1^fl/fl^ and HMGB1^ΔHep^ mice after a fasting/refeeding challenge. Data are means ± SEM from n=7 (HMGB1^fl/fl^) or n=8 (HMGB1^ΔHep^) per group for the HFD protocol (**A-C**) and from n=8 (HMGB1^fl/fl^) or n=8 (HMGB1^ΔHep^) per group for the F/R protocol (**D-F**). *p<0.05, **p<0.01, ***p<0.001, ****p<0.0001 by unpaired Mann and Whitney comparison.

### Nuclear HMGB1 represses hepatocyte lipogenesis in vivo and in vitro in a cell-autonomous manner

The enhanced hepatosteatosis in HMGB1ΔHep mice may result from an increased activity of lipogenesis in the hepatocytes. To address this question, hepatic lipid synthesis was monitored *in vivo* using radiolabeled substrates upon a fasting-refeeding challenge (Fig. 2A). After 6 hours of fasting, HMGB1fl/fl and HMGB1ΔHep mice received a bolus of 3H glucose, and the 3H radioisotope incorporation was quantified in the lipid fractions of several tissues after 8 hours of refeeding. Upon CD, while F/R induced a strong 3H incorporation mainly in brown adipose tissue (BAT) and liver of HMGB1fl/fl mice (Fig. 2A), this effect was even more pronounced in HMGB1ΔHep mice, suggesting a higher capacity of *Hmgb1-null* hepatocytes to synthesize lipids after refeeding (Fig. 2A). In parallel, we evaluated *in vivo*, a potential disturbance of lipoprotein metabolism in HMGB1ΔHep mice upon CD and HFD. The VLDL secretion after treatment with the lipoprotein lipase inhibitor tyloxapol (Fig. 2B) and the activity of the microsomal triglyceride transfer protein (MTP), a key enzyme involved in lipid export (Fig. 2C), were both identical in HMGB1fl/fl and HMGB1ΔHep mice subjected to CD and HFD.

**Fig. 2.**
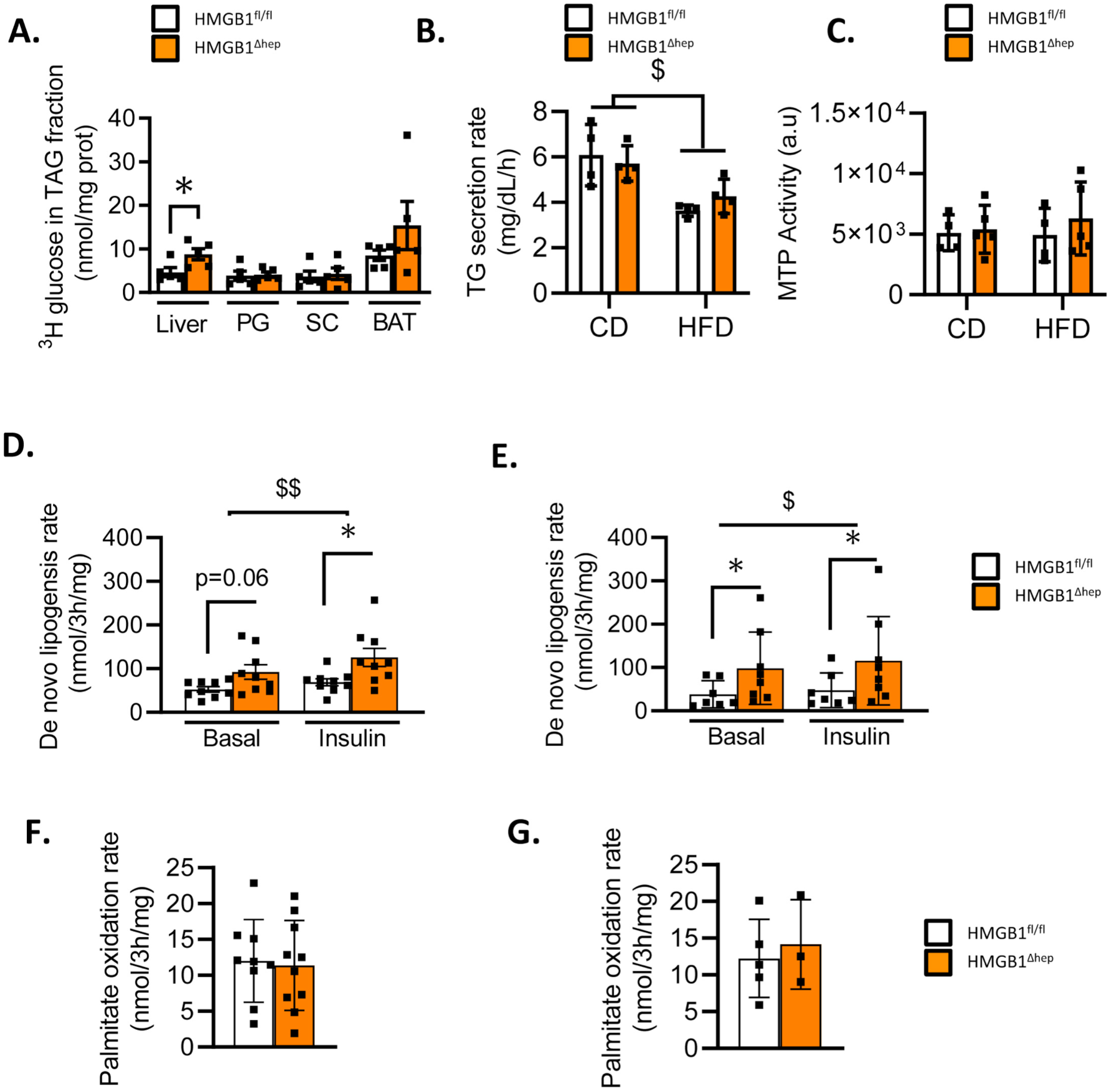
*Hmgb1* deletion increases hepatocyte lipid synthesis *in vitro* and *in vivo*. (**A**) *In vivo*, lipogenesis was measured on HMGB1^fl/fl^ (n=5) and HMGB1^ΔHep^ (n=5) mice. Mice were food deprived for six hours then injected with ^3^H glucose (0.4 μCi/g, i.p) and euthanized one hour later and ^3^H was measured in TAG fraction of liver, adipose tissues (PG, SC and BAT). (**B**-**C**) *In vivo*, assessment of liver lipoprotein secretions determined by (**B**) measuring circulating tri-acyl glycerol concentration (n=4 per genotype and diet) and (**C**) liver MTP activity, HMGB1^fl/fl^ (n=4) and HMGB1^ΔHep^ (n=5). (**D-E**) Lipid synthesis was measured *in vitro*, on primary hepatocytes isolated from adult HMGB1^fl/fl^ (n=7-9) and HMGB1^ΔHep^ (n=8-9) mice on (**D**) chow diet and (**E**) HFD. (**F-G**) Beta-oxidation was measured *in vitro*, on primary hepatocytes isolated from adult HMGB1^fl/fl^ and HMGB1^ΔHep^ mice on (**F**) chow diet (HMGB1^fl/fl^ n=9 and HMGB1^ΔHep^ n=10) and (**G**) HFD (HMGB1^fl/fl^ n=5 and HMGB1^ΔHep^ n=3). Data are means ± SEM of three independent experiments. p<0.05, **p<0.01, ***p<0.001, ****p<0.0001 by unpaired Mann and Whitney comparison or two-way ANOVA. $ p<0.05, $$ p<0.01, $$$ p<0.001, for treatment effect by one-way ANOVA.

Present knowledge indicates that the regulation of hepatic lipogenesis depends on the interplay, within the liver, between hepatocytes and non-parenchymal cells and is also influenced by other tissues, mainly the adipose tissue. Therefore, we interrogated whether the increase of liver lipogenesis in HMGB1^ΔHep^ mice could be cell-autonomous. To address this point, primary hepatocytes were isolated from HMGB1^fl/fl^ and HMGB1^ΔHep^ mice and lipogenic activity was assessed *in vitro* using the same strategy as described above for the *in vivo* study. Consistent with the *in vivo* data, after isolation from mice under CD, cultured HMGB1^ΔHep^ hepatocytes displayed an increased lipogenic activity compared to HMGB1^fl/fl^ hepatocytes (**Fig. 2D**).

However, lipogenesis was stimulated to the same extent by insulin (**Fig. 2D**) in hepatocytes from both genotypes. Interestingly, when isolated from HFD-fed mice, HMGB1^ΔHep^ hepatocytes still exhibited a higher lipogenic activity compared to HMGB1^fl/fl^ hepatocytes (**Fig. 2E**) and insulin slightly increased the lipogenesis independently of the genotypes. Importantly, palmitate oxidation was also measured in primary hepatocytes from both genotypes, and no difference in lipid utilization was observed neither upon CD (**Fig. 2F**) nor HFD (**Fig. 2G**). Collectively, these results suggest that HMGB1 represses lipogenesis in hepatocytes in a cell autonomous-manner, without affecting FA oxidation.

### Hepatic deletion of Hmgb1 affects specifically liver insulin sensitivity

Studies have reported a strong correlation between hepatic lipid accumulation and a decreased insulin sensitivity in the liver (*30*). Therefore, we next monitored whether the liver steatosis induced by hepatocyte *Hmgb1* deletion has any effect on glucose homeostasis and/or insulin signaling in mice subjected to a HFD60%. Upon HFD both HMGB1^fl/fl^ and HMGB1^ΔHep^, displayed a similar glucose homeostasis and global insulin sensitivity (**Fig. 3A-C**), albeit a slight trend toward a higher AUC after oral glucose test tolerance was observed in HMGB1^ΔHep^ mice (**Fig. 3A**). Of note, insulin levels either after starvation or after a bolus of glucose were similar between both groups (**Fig. 3B**), ruling out that hepatic *Hmgb1* deletion may interfere with insulin secretion. Interestingly HMGB1^ΔHep^ mice displayed a higher glycaemia after 14 hours starvation (**Fig. 3D**), corroborated by a higher AUC during a pyruvate tolerance test compared to HMGB1^fl/fl^ mice (**Fig. 3E**), suggesting an increased hepatic glucose production consistent with a potential hepatic insulin resistance. In addition, liver glycogen content was lower in HMGB1^ΔHep^ mice as shown by the PAS coloration (**Fig. 3F-G**) supporting a compromised glycogen synthesis. All together these data show that the increased hepatosteatosis in HMGB1^ΔHep^ mice is associated with a noticeable perturbation of insulin signaling. This was confirmed by the lower level of AKT phosphorylation, recognized as a classic downstream effector of the insulin receptor, in the liver of HMGB1^ΔHep^ mice subjected to a 12 week-HFD, compared to HMGB1^fl/fl^ mice (**Fig. 3H**). To functionally test a possible alteration of insulin sensitivity in absence of hepatocyte *Hmgb1*, HMGB1^fl/fl^ and HMGB1^ΔHep^ mice subjected to CD or a long term HFD (24 weeks-as the global insulin signaling is more perturbed compared to a 12 week-HFD) were challenged with an acute injection of insulin (0.75U/kg) or saline (**Fig. 3I**). In CD-fed mice of both genotypes, we observed no differences in the insulin-induced phosphorylation of AKT, compared to saline conditions (fig. S2F). In HFD-fed mice, insulin injection induced the expected phosphorylation of AKT in the liver, adipose tissue and skeletal muscle (**Fig. 3I**) in control mice, but remarkably the amount of p-AKT was much lower selectively in liver samples harvested from HMGB1^ΔHep^ mice, compared to skeletal muscle and adipose tissue (**Fig. 3I**). Collectively these data show a selective impact of hepatocellular HMGB1 deficiency on liver insulin signaling upon long term-HFD feeding.

**Fig. 3.**
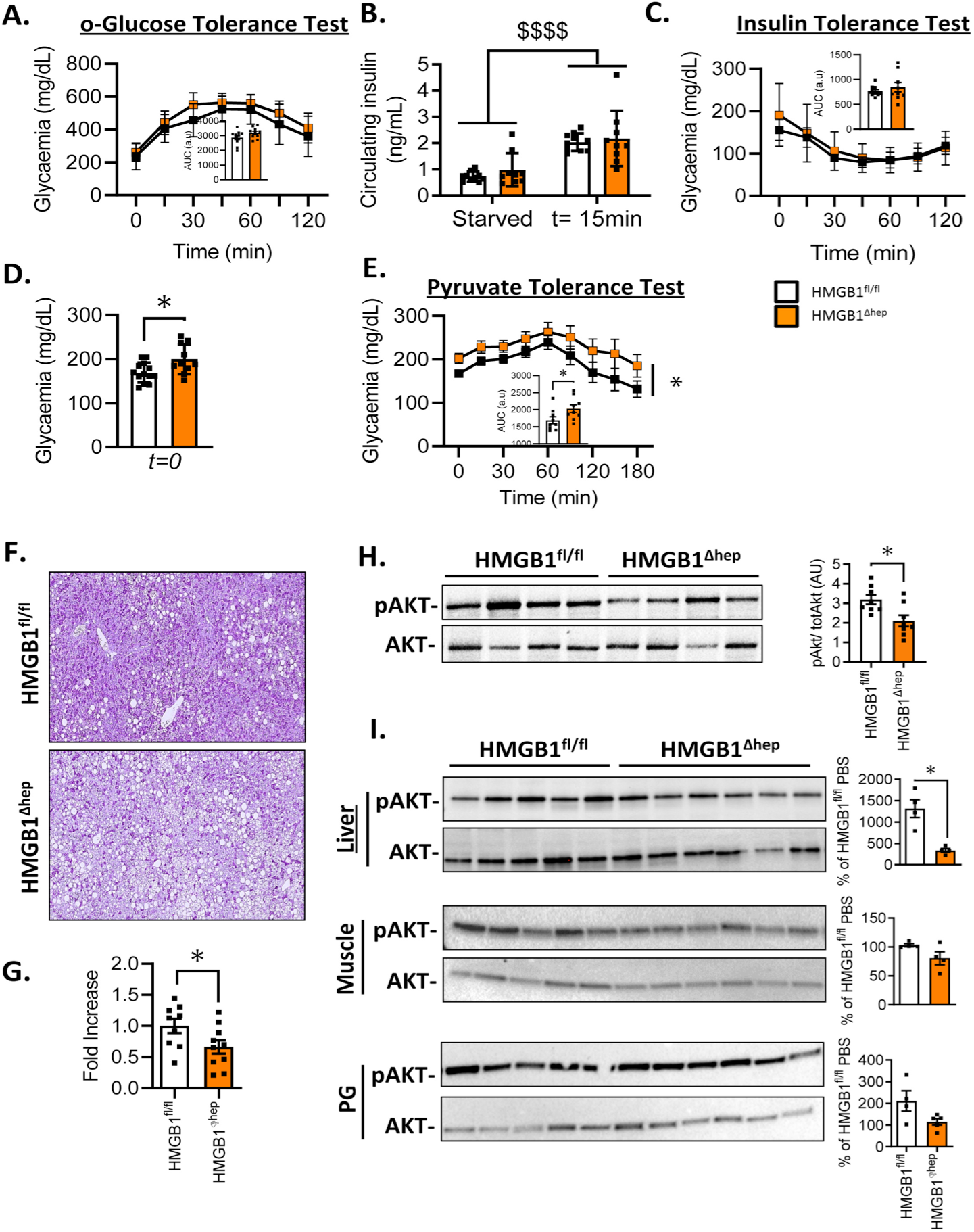
Hepatocyte specific *Hmgb1* deleted mice on HFD display reduced insulin sensitivity in the liver. (**A**) Analysis of oral glucose tolerance test, (**B**) Insulin levels after fasting or 15 minutes post glucose bolus, (**C**) insulin tolerance test (**D**) Fasting glycaemia levels after 16 hours of fasting. and (**E**) pyruvate tolerance test on HMGB1^fl/fl^ and HMGB1^ΔHep^ mice fed on HFD for 12 weeks. (**F**) Hepatic PAS staining representative images with (**G**) quantification on HMGB1^fl/fl^ and HMGB1^ΔHep^ mice fed on HFD for 12 weeks. (**H**) Representative immunoblot targeting p-AKT and tot-AKT with quantification performed on the whole animal cohort, on liver biopsies from on HMGB1^fl/fl^ and HMGB1^ΔHep^ mice fed on HFD for 12 weeks. (**I**) Representative immunoblot targeting p-AKT and tot-AKT with quantification performed on the whole animal cohort, on liver biopsies from HMGB1^fl/fl^ and HMGB1^ΔHep^ mice fed on HFD for 24 weeks, starved 4 hours and injected with insulin (i.p. 0.75U/kg-15 minutes). Data are means ± SEM from n=10 (HMGB1^fl/fl^) or n=11 (HMGB1^ΔHep^) per group for the HFD protocol (**A-H**) and from n=4 (HMGB1^fl/fl^) or n=4 (HMGB1^ΔHep^) per group for the HFD 24-week with acute injection of insulin protocol (**I**). *p<0.05, **p<0.01, ***p<0.001, ****p<0,0001 by unpaired Mann and Whitney comparison or two-way ANOVA.

### The signaling of LXR is enhanced in the absence of Hmgb1

To unveil the signaling pathways regulated by HMGB1, we performed gene expression profiling using cDNA microarray of HMGB1^fl/fl^ and HMGB1^ΔHep^ liver samples from mice subjected to a 12 week-HFD regimen or a F/R challenge (**Fig. 4**). Microarray analysis and unsupervised clustering displayed on the heatmaps showed that deletion of *Hmgb1* caused significant changes in the liver transcriptome (**Fig. 4A**, fig. S4A-B). Venn diagrams revealed that in liver samples from HMGB1^ΔHep^ mice, there were 295 up- and 471 down-regulated genes upon HFD and 125 up- and 380 down-regulated genes after F/R (**Fig. 4B**). Of note, as displayed in the Venn diagram (**Fig. 4B**), 253 genes (roughly 25%) of the identified genes are similarly regulated in both challenges (HFD and F/R). Hierarchical clustering method showed that the vast majority of these genes are subjected to the same type of variations in both conditions (**Fig. 4C**) suggesting that these groups of genes belong to pathways under robust regulation by *Hmgb1*. The enrichment analysis of these 253 common genes, using the EnrichR database, indicated that among all gene ontology (GO) terms represented in HMGB1^ΔHep^ livers, the most enriched GO terms were “metabolism of lipids” and “metabolism” (**Fig. 4D-E**), confirming our histological findings. Based on the analysis of the gene network using the Reactome database, numerous genes regulated by HMGB1 in both nutritional conditions, are connected to metabolism functions, and more specifically, to lipid metabolism (**Fig. 4F**). We then narrowed our focus on gene clusters involved in these identified GO terms, and further performed analysis on potential upstream regulators involved, by using EnrichR database (**Fig. 4G**). Interestingly, among the identified transcription factors, LXR and PPARγ came up with the highest score. LXRα and PPARγ are well known for their critical pro-lipogenic activity in the liver, which is in line with the phenotype displayed by the HMGB1^ΔHep^ mice (**Fig. 1**).

**Fig. 4.**
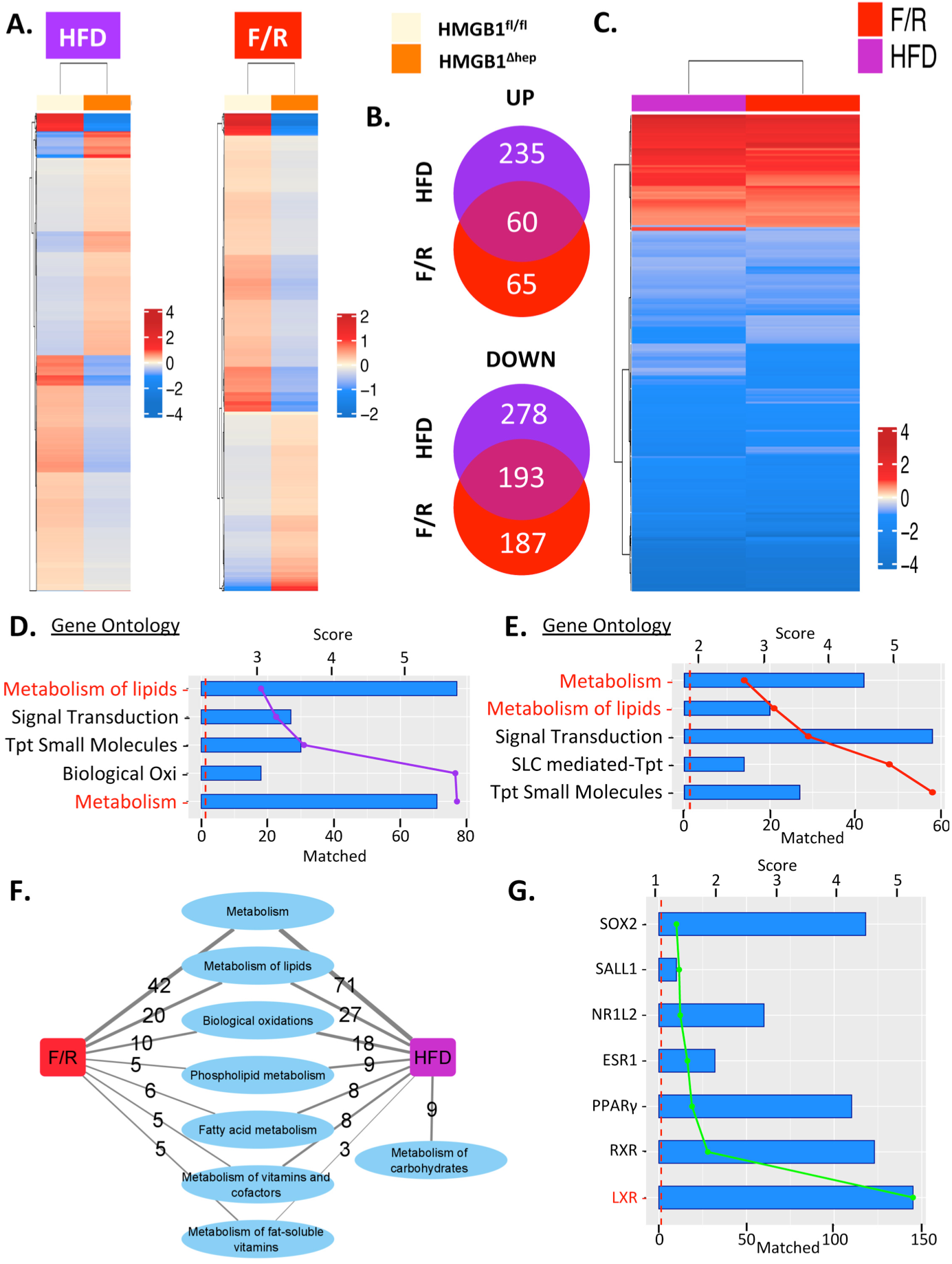
Microarray analysis of hepatic gene expression profiles in HMGB1^ΔHep^ mice. (**A**) Heatmap showing genes that are differentially expressed in the livers of HMGB1^ΔHep^ mice compared to HMGB1^fl/fl^ mice (fold change > 1.5; P-Value <= 0.01) after HFD (left panel) or F/R (right panel). Heatmaps display the mean normalized expression per genotype per nutritional challenge. (**B**) Venn Diagram displaying overlap between up and down regulated genes in the two regimens. (**C**) Heatmap displaying only DEG commonly found in both regimens (fold change > 1.5; P-Value <= 0.01). (**D-E**) Top 5 GO biological processes enriched using gene sets for each regimen, with the −log10(P-Value) of enrichment shown as bars and the number of matched genes as colored lines. (**F**) Network displaying Reactome pathways related to metabolism that are enriched by our HMGB1 gene sets from both nutritional challenges, edge thickness represents the number of genes regulated by HMGB1 among each sub-category. (**G**) Top upstream regulators identified using the ChEA database, with the −log10(P-Value) of enrichment as bars and the number of gene matched per as a green line. Data are means ± SEM from n=4 (HMGB1^fl/fl^) or n=4 (HMGB1^ΔHep^) per group for the 12 week-HFD protocol and from n=4 (HMGB1^fl/fl^) or n=4 (HMGB1^ΔHep^) per group for the F/R protocol.

Collectively our unbiased transcriptomic study indicated that in the liver upon metabolic stress, HMGB1 might repress the expression of gene clusters partly controlled by LXRα and PPARγ and involved in hepatic lipid synthesis.

### Exaggerated hepatic steatosis in the Hmgb1-null liver is dependent on LXRα activity

As LXRα is a key lipogenic transcription factor involved in cholesterol metabolism and liver lipogenesis, the potential de-repression of its activity induced by HMGB1 deletion could translates into liver steatosis (*31*, *32*). However, it is less clear whether PPARγ is a significant trigger of liver steatosis. The role of PPARγ in HFD-induced hepatosteatosis is supported by several reports (*33*, *34*), but no studies have investigated its potential role during F/R-induced liver steatosis. To clarify this, we subjected mice carrying hepatocyte specific-*Pparγ* deletion to F/R challenge, and the results show no major contribution of hepatocyte PPARγ to the progression of F/R-induced liver steatosis (fig. S5) based on liver body weight ratio, Oil Red-O staining, neutral lipid profile or mRNA expression of hepatic steatosis markers (fig. S5A-D). This suggests that PPARγ, *per se*, is not a determinant trigger of hepatic lipogenesis, and therefore its potential contribution in the severe steatosis displayed in HMGB1^ΔHep^ mice is likely minor.

Subsequently, we focused on the functional interdependence between HMGB1 and LXRα, examining the effect of pharmacological activation and adenovirally-mediated inhibition of LXRα in HMGB1^fl/fl^ and HMGB1^ΔHep^ mice (**Fig. 5**). To establish a possible causal link between the absence of HMGB1 and LXRα activity, the HMGB1^fl/fl^ and HMGB1^ΔHep^ mice were treated with a synthetic LXR agonist (T0901317) for four consecutive days (30mg/kg-*per os*) (**Fig. 5A-B**, fig. S6A). Remarkably, already before treatment, several LXRα dependent genes (*Srebf1*, *Fasn, Elovl-6*, *Abcg5*, and *Abcg-8*) were up-regulated in the HMGB1^ΔHep^ livers (**Fig. 5A**). As expected, T0901317 treatment of HMGB1^fl/fl^ mice potently induced expression of LXR dependent genes (*Srebf1*, *Fasn, Elovl-6*, *Scd-1*, *Abcg5*, *Abcg-8*) in the liver compared to vehicle treated HMGB1^fl/fl^ mice. Importantly, HMGB1^ΔHep^ livers displayed a significantly higher response to T0901317 than HMGB1^fl/fl^ mice, with an enhanced expression of *Fasn*, *Elovl-6*, *Abcg-5* and *Abcg-8* (**Fig. 5A** and fig. S6A). This higher response was corroborated by histological examination showing an increased Oil Red-O staining in *Hmgb1* deleted livers in mice subjected to the T0901317 treatment (**Fig. 5B**). Taken together these results indicate that the higher lipogenesis in HMGB1^ΔHep^ livers is likely due to an enhanced LXRα activity. To complement this study, and firmly establish the role of LXRα in the enhanced hepatic steatosis seen in HMGB1^ΔHep^ mice, we silenced LXRα expression *in vivo*, using an adenovirus expressing shRNA targeting the receptor (Ad-Sh*Lxrα*) (**Fig. 5C-D** and fig. S6B-C). Seven days after viral infection, the hepatic LXRα, but not β, mRNA levels were reduced showing that expression of LXRα, as long as LXRα -dependent genes, were successfully blunted in Ad-Sh*Lxrα* injected animals compared to control animals injected with an adenovirus expressing a scrambled shRNA (Ad-Sh*SCR*), highlighting the potency and specificity of LXRα targeting (fig. S6B-C). Consistent with the results presented above, Ad-sh*SCR* treated HMGB1^ΔHep^ mice displayed increased hepatic steatosis compared to Ad-sh*SCR* injected HMGB1^fl/fl^ mice either upon F/R (**Fig. 5C**) or HFD feeding (**Fig. 5D**) as shown by Oil Red-O staining. And remarkably, Ad-sh*Lxrα* treatment lowered drastically hepatic steatosis in both groups of animals (**Fig. 5C-D**), suggesting that LXRα plays a major role in the enhanced hepatic lipid synthesis of HMGB1^ΔHep^ mice. In summary, these results suggest that the LXRα activity is responsible for the enhanced hepatic lipid synthesis in *Hmgb1*-null livers and support a repressive role of HMGB1 on hepatic lipogenesis through repression of LXRα activity.

**Fig. 5.**
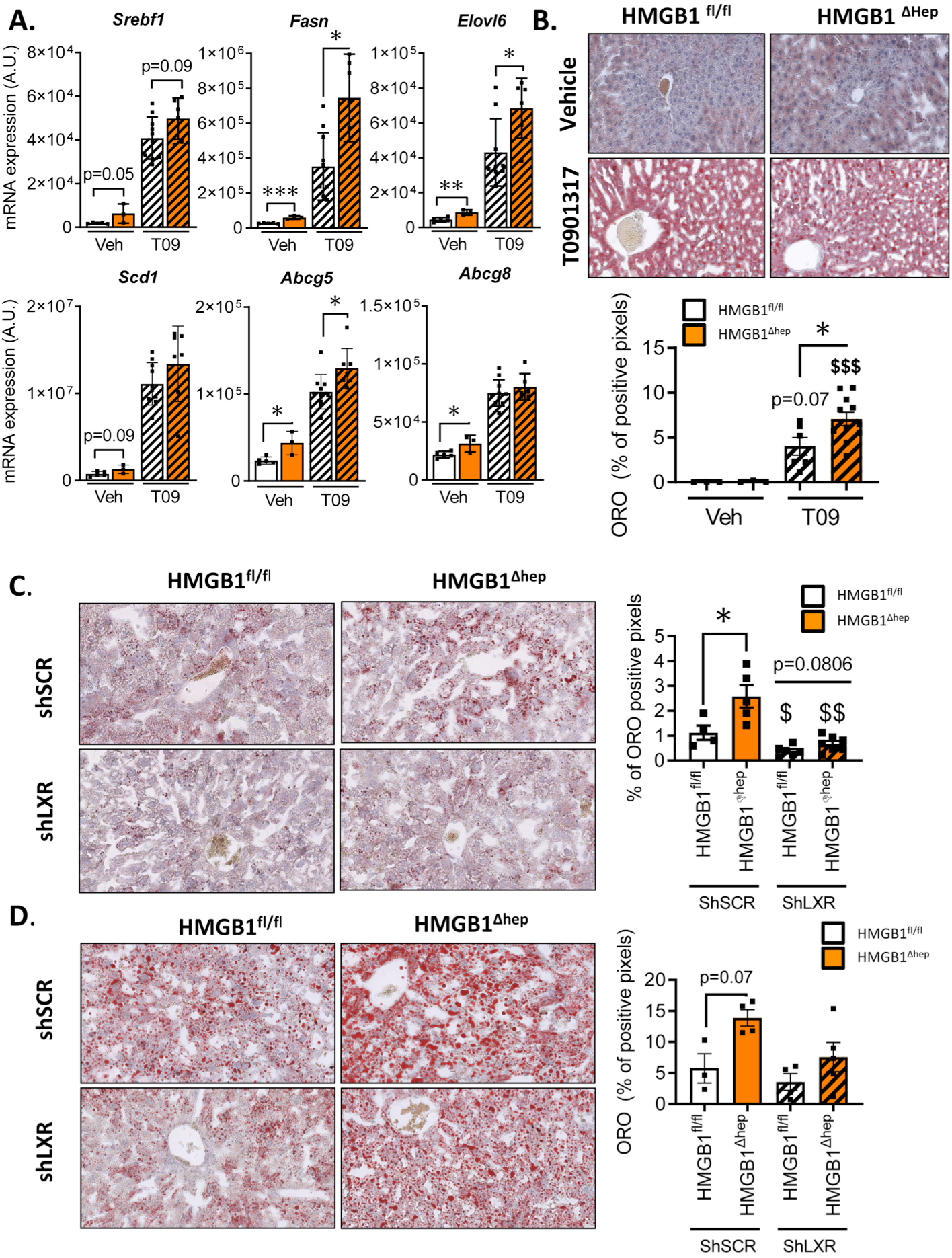
*In vivo* knockdown of LXR normalizes liver steatosis in HMGB1^ΔHep^ mice. (**A-B**) HMGB1^fl/fl^ (n=15) and HMGB1^ΔHep^ (n=9) mice were treated either with vehicle (5% carboxy-methyl-cellulose) or LXR synthetic agonist T0901317 (oral gavage, 30 mg/kg/day) for four consecutive days, after 6 hours starvation on the last day mice were sacrificed. (**A**) Liver tissue was then subjected to RT-qPCR analysis of the indicated LXR dependent genes and (**B**) liver steatosis was quantified using Oil Red-O staining. (**C**) HMGB1^fl/fl^ (n=10) and HMGB1^ΔHep^ (n=12) mice were infected with either adenovirus expressing a LXR shRNA or a scramble (SCR) sequence, then subjected 7 days later to a F/R challenge. Liver steatosis was determined by Oil Red-O staining on liver sections with the quantitative representation displayed on the right. (**D**) HMGB1^fl/fl^ and HMGB1^ΔHep^ mice were subjected to HFD for four weeks and then infected with either adenovirus expressing a LXRα shRNA (n=7) or a scramble shRNA (SCR) n=9) sequence and euthanized 7 days later. Liver steatosis was assessed by Oil Red-O staining on liver section tissue with the quantitative representation displayed on the right. Data are means ± SEM. *p<0.05, **p<0.01, ***p<0.001, ****p<0.0001 HMGB1^fl/fl^ and HMGB1^ΔHep^ comparison, by unpaired Mann and Whitney comparison. $ p<0.05, $$ p<0.01, $$$ p<0.001, for treatment effect by one-way ANOVA.

### HMGB1 binds to LXRα target genes involved in lipogenesis

Having identified LXRα as potential targets for repression by HMGB1, we determined the molecular mechanisms by which HMGB1 is exerting this action. Considering the impact HMGB1 may have on chromatin compaction (*22*), we first performed an assay for transposase-accessible chromatin using high throughput sequencing (ATAC-seq) to evaluate the global chromatin dynamics in the absence of hepatic-HMGB1. Hepatocyte nuclei were purified from liver samples harvested from HMGB1^fl/fl^ and HMGB1^ΔHep^ mice upon CD feeding or after FR (fig. S7). Remarkably, at basal state the principal component analysis (PCA) analysis of the ATAC-seq peaks revealed no distinct pattern in chromatin states between both genotypes (fig. S7A), in reads alignment in a genome browser (fig. S7B) or in the open chromatin regions (OCR) locations around transcription start sites (TSS) (fig. S7C). In sharp contrast, F/R in HMGB1^fl/fl^ mice triggered significant changes in chromatin state compared to the CD condition (respectively 68776 vs. 47725 OCRs), but similar modifications were detected in the liver chromatin from F/R HMGB1^ΔHep^ mice. Strikingly, only 4 OCRs were differentially nucleosome-depleted between both genotypes supported by the very high number of common aligned peaks (fig. S7D). PCA analysis, examination of TSS charts and annotation chart-pie confirmed the high similarity in the chromatin state of both librairies (fig. S7E-G). A close visualization of aligned peaks in loci of lipogenic genes regulated by LXRα (*Srebf1*, *Scd-1*, *Cidec* or *Fasn*) (fig. S7H) showed as expected the same chromatin state pattern between both genotypes. As presumed from this very low number of sites differentially opened in the chromatin between control and *Hmgb1* null-livers, enrichment analysis could not identify any statistically significant biological functions related to these modifications. Overall the analysis of ATAC-seq datasets ruled out a putative model where HMGB1 may regulate hepatic lipid metabolism through chromatin packaging.

Next we sought to determine, using chromatin immuno-precipitation combined with high-throughput sequencing (ChIP-sequencing), whether HMGB1 might exert its activity on gene transcription directly through its abilities to bind DNA. We first set up a reliable and robust ChIP protocol on cells *in vitro*, as HMGB1 ChIPing might be challenging (*26*) (fig. S8A-C). Then, using frozen liver samples, we examined HMGB1 binding genome-wide in HMGB1^fl/fl^ under CD, HFD, and after F/R (**Fig.6** and fig. S8-S9). Of note, HMGB1 ChIP-seq was also performed on HMGB1^ΔHep^ livers and these datasets were used as negative control to determine non-specific signals. These background peaks were subtracted in libraries from HMGB1^fl/fl^ livers (fig. S8D-F). Under CD feeding condition, 201250 peaks were detected on the whole genome that were predominantly located in promoters (18.5%), introns (29.3%) and intergenic regions (32.6%) (fig. S9A). Interestingly only 155854 and 32006 peaks were detected under the F/R or HFD conditions, respectively, suggesting a significant remodeling of the HMGB1 binding pattern during metabolic stress, even though the qualitative binding remains nearly the same (fig. S9A). The PCA plot of **Figure 6A** demonstrates significant global differences in HMGB1 DNA occupancy between CD versus F/R and HFD. Venn diagram confirmed this trend with only a few peaks (8859) detected in common in the three conditions (**Fig. 6B**). The genome browser view of chromosome 3, 12 and 14 exemplified the drastic repositioning of HMGB1 upon nutritional stress (**Fig. 6C**). Along the same lines of observation, partitioning of HMGB1-bound sites by distance to TSSs confirmed the severe change in DNA occupancy of HMGB1. Importantly, the results suggested that most HMGB1 sites located around the TSSs (+/− 3000 bp) under CD feeding were not used under the F/R or HFD conditions (**Fig. 6D**). Enrichment analysis based on peaks differentially called in CD vs HFD feeding (**Fig. 6E**) and CD vs F/R (**Fig. 6F**) revealed that among several biological functions (GO categories), two are remarkably related to lipid metabolism as the “integration of energy metabolism” and “phospholipid metabolism” **(Fig. 6E-F**). In these two GO categories, 134 genes displayed a very high occupation rate upon CD compared to F/R and HFD and nearly 90% of these genes displayed a lower occupancy of HMGB1 in both challenges when compared to CD. These results suggest a common mechanism of regulation in F/R and HFD (**Fig. 6G**, full list in **Table S1 and Table S2**). To gain insight into the gene expression program regulated by HMGB1, we performed a motif identification analysis on 134 genes unveiled by the enrichment analysis. The oPOSSUM-3 motif tool revealed the binding motifs of the transcription factors of LXR, identifying this nuclear receptor among the top regulators (**Fig. 6H**). To functionally test whether the HMGB1 occupancy rate would have an incidence on the level of gene expression, we went back to the microarray data to measure the expression of the 134 genes identified in the enrichment analysis performed above. Out of the 134 genes, 70 and 78 are up-regulated in HFD and F/R, respectively, in livers from HMGB1^ΔHep^ mice compared to HMGB1^fl/fl^ mice (**Fig. 6I-J**) providing evidence for a negative correlation between the HMGB1 DNA occupation and the expression of metabolic related-genes identified in the ChIPseq. These data demonstrate that HMGB1 may play a suppressive action on LXRα activity and, consequently, on the level of expression of its target genes.

**Fig. 6.**
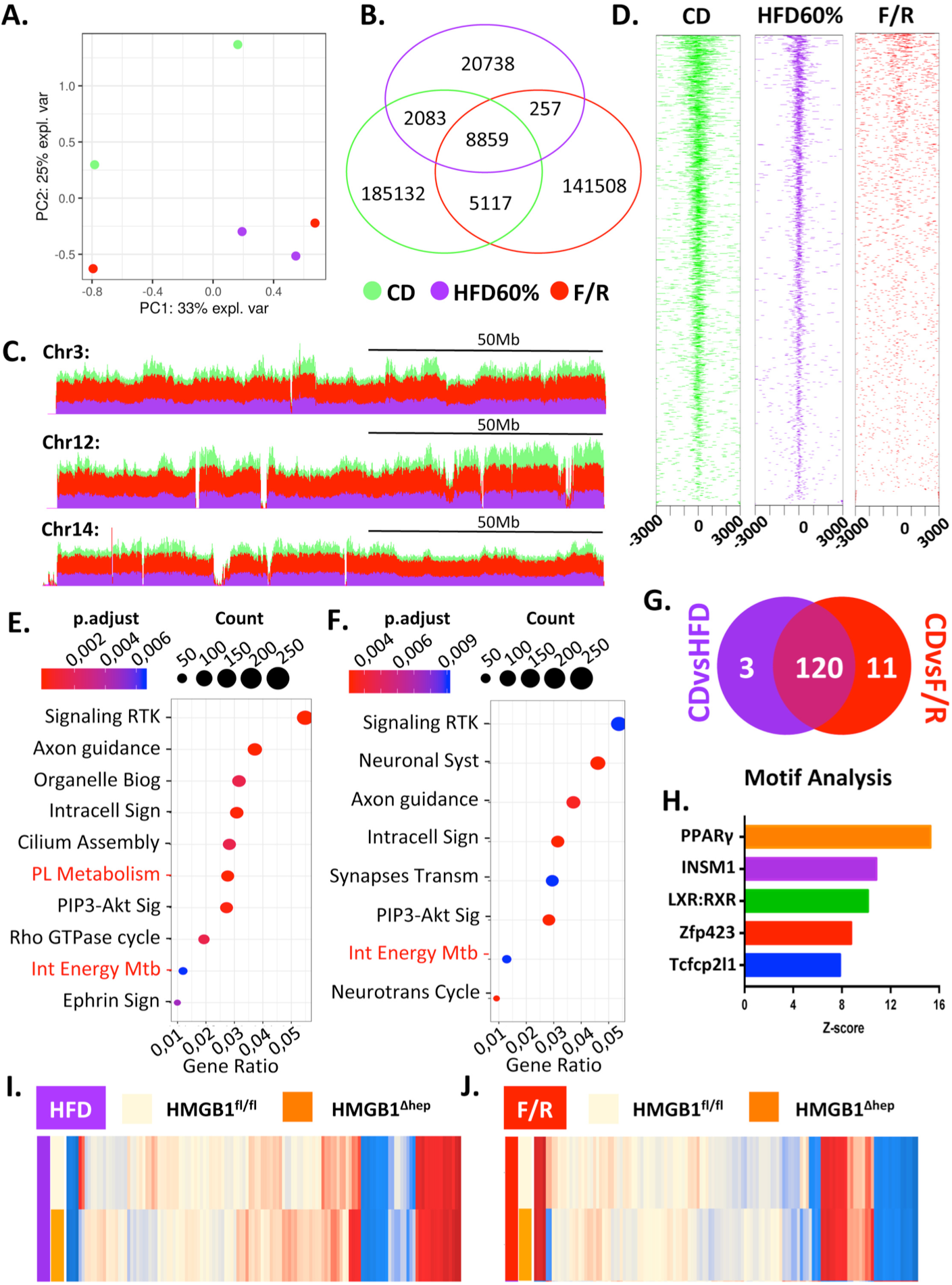
ChIP-seq identified a subset of LXR responsive genes to be negatively regulated by HMGB1 during liver steatosis. (**A**) Principal component analysis scores plot of ChIP-seq data of liver tissue from HMGB1^fl/fl^ mice on chow diet (green) or subjected to F/R (red) or HFD (purple). (**B**) Venn Diagram showing the number of HMGB1 binding peaks, (**C**) UCSC genome browser of tracks (stacked) showing HMGB1 differential chromatin occupancy and (**D**) average signal density profiles around transcription starting site in different nutritional states: chow diet (green) or during HFD (purple) or after F/R (red). (**E-F**) Functional enrichment analyses showing GO terms associated with the differential HMGB1chromatin binding sites between (**E**) chow diet and HFD and (**F**) chow diet and F/R. (**G**) Venn Diagram displaying shared enriched genes (n=134) displaying a very high occupancy rate during fed state belonging to “Integration of energy metabolism” and “Phospholipid metabolism” GO functions compared to HFD (purple) and F/R (red). (**H**) Graph bar displaying consensus motifs in promoters of the 134 genes differentially occupied by HMGB1 via OPOSUM analysis; the bars represent the z-score. (**I-J**) Heatmaps displaying the mean microarray expression levels for the 134 genes identified by ChIP-seq in liver from HMGB1^fl/fl^ (n=4) and HMGB1^ΔHep^ (n=4) mice subjected to either HFD (**I**) or F/R (**J**).

Taken together, our data are in support of a model whereby at basal state (CD), HMGB1 binds to chromatin loci to modulate the transcription of a number of genes controlled by LXRα which are particularly involved in energy metabolism and lipogenesis.

### In vitro, HMGB1 exerts a repressive action on LXRα

Since HMGB1 modulates chromatin structure and, therefore, regulates transcription factor activity, we examined whether HMGB1 could inhibit LXRα transcriptional activation in cultured cells transfected with luciferase reporter genes harboring LXR response elements (LXRE). Expression of HMGB1 dramatically decreased LXRα transcriptional activity already at basal state but also after pharmacological activation by synthetic LXR (T093911) or RXR (LG268) agonists (**Fig. 7A**). Next, we tested whether HMGB1 directly interacts with LXRα *in vitro* co-immunoprecipitation assays (**Fig. 7B**) but no interaction could be detected between a flagged Myc-HMGB1 and HA-LXRα (**Fig. 7B**). These *in vitro* assays help to firmly establish that HMGB1 is capable of potently repress LXRα activity at basal state but also upon pharmacological activation, but without any direct physical interaction. Therefore, we tested whether HMGB1 mediated-inhibition of LXR activity may occur through suppressing LXR interaction with the DNA encoding LXR target genes. The ChIP sequencing data suggested that the localization of HMGB1 at specific gene loci correlated with its repressive role of LXR target genes such as *Acly* or *Fasn*. These two loci were significantly enriched in CD (green tracks) compared to HFD (purple tracks) and F/R (red tracks) (**Fig. 7C-D**). Interestingly, HMGB1 bound across the whole loci (**Fig. 7C-D)** and the promoters of the two HMGB1 repressed genes, *Acly* or *Fasn* displayed a heterogeneous HMGB1 occupation patterns (fig. S9B-C), with *Acly* promoter displaying a high occupation rate in the TSS as opposed to *Fasn* promoter (fig. S9B-C). This suggests that HMGB1 is not exerting its repressive effect only through TSS occupation. This prompted us to extend the analysis to a series of key genes involved in lipogenesis by performing RT-qPCR experiments on liver samples from adult HMGB1^fl/fl^ and HMGB1^ΔHep^ mice fed with CD, a condition under which HMGB1 repression was strong. The results showed a consistent up-regulation in the expression level of key lipogenic genes when HMGB1 was lacking in livers of HMGB1^ΔHep^ mice. The expression of direct LXRα target genes such as *Srebf1*, *Scd-1*, *Abcg-5* or *Abcg-8* and indirect target genes such as *Cd36*, *Cidec*, *Pnpla3* or *Fasn* (**Fig. 7E**) was increased in the liver of these mice compared to their floxed littermates. To establish a causal link between the nuclear presence of HMGB1 and the mRNA expression level of the above-mentioned genes, we deleted HMGB1 selectively in hepatocytes using the hepatocyte-specific promoter of the thyroxine-binding globulin (TBG) gene to express the Cre-recombinase via an AAV8-vector (AAV8-TBG-Cre) in adult HMGB1^fl/fl^ mice. This strategy was validated by the lower levels of HMGB1 mRNA and protein levels detected in the liver of AAV8-TBG-Cre expressing mice compared to the control group (fig. S10A-B). Remarkably seven days post viral infection with the recombinant virus, the reduced *Hmgb1* expression resulted in up-regulation of a vast majority of LXRα responsive genes, similarly to what is seen in liver of mice with a constitutive *Hmgb1* deletion in hepatocytes (**Fig. 7F**) This result supports a causal and repressive role for HMGB1 on the level of expression of this subset of genes. Overall, these findings support that HMGB1 is repressing LXRα transcriptional activity, which is not mediated by a direct physical interaction with the receptor but rather through a complex DNA occupation across the LXRα responsive gene loci.

**Fig. 7.**
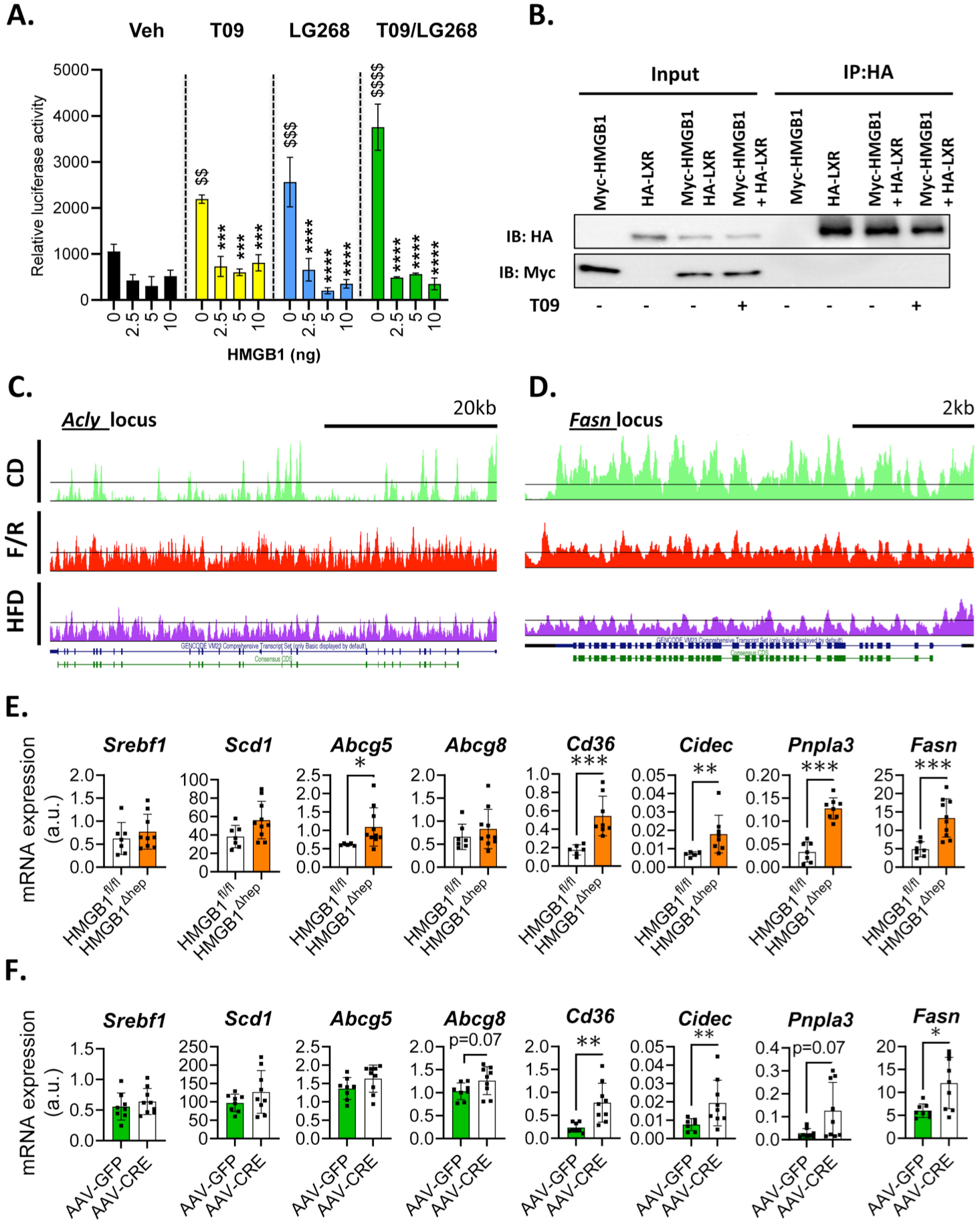
HMGB1 represses LXRα transcriptional activity *in vitro*. (**A**) Effect of HMGB1 on LXRE-luciferase reporter activity. Ad293 cells were treated with DMSO (vehicle), T0901317 (noted T09) (0.1 uM) and/or LG286 (1 nM) for 14 hours. (**B**) Co-immunoprecipitation assay was performed to detect a potential interaction between HMGB1 and LXR in Ad293 transfected cells treated with DMSO (vehicle) or T0901317 (0.1 nM for 14 hours. Data are representative of three independent experiments. (**C-D**) Genome browser shot of ChIP-seq data along the locus of *Acly* and *Fasn* gene loci in liver from HMGB1^fl/fl^ and HMGB1^ΔHep^ mice upon chow diet (green), HFD (purple) and after F/R (red). Gene (blue) and CDS (green) models are displayed on the bottom track. (**E**) Gene expression of direct (*Srebf1*, *Scd-1*, *Abcg-5* and *Abcg-8*) and indirect (*Cd-36*, *Cidec*, *Pnpla3* and *Fasn*) targets of LXRα in livers of HMGB1^fl/fl^ (n=7) and HMGB1^ΔHep^ (n=9) mice. (**F**) Adult HMGB1^fl/fl^ mice were infected either with AAV8-Gfp (n=8) or AAV8-TBG-Cre (n=9) to selectively generate *Hmgb1* deletion in hepatocytes *in vivo* and expression of direct (*Srebf1*, *Scd-1*, *Abcg-5* and *Abcg-8*) and indirect (*Cd-36*, *Cidec*, *Pnpla3* and *Fasn*) responsive genes were determined using RT-qPCR. Data are means ± SEM of three independent experiments. *p<0.05, **p<0.01, ***p<0.001, ****p<0.0001 by unpaired Mann and Whitney comparison or two-way ANOVA. **$** p<0.05, **$$** p<0.01, **$$$** p<0.001, for treatment effect by two-way ANOVA.

## Discussion

Lipogenesis is a fundamental function of the liver to regulate and buffer the amount of circulating lipids, which could present a risk of cellular toxicity in the long run, for numerous tissues (*35*). Hepatic lipogenesis is therefore tightly regulated by a large number of factors, including TFs and nuclear proteins that together manage positive and repressive actions on gene transcription. These regulatory processes and their interplay are complex and only partly understood and have high relevance due to the high world-wide prevalence of NAFLD (*1*). Herein, we unraveled a new mechanism regulating liver lipogenesis involving the nuclear factor HMGB1. Using both constitutive and induced knockouts of *Hmgb1* gene selectively in hepatocytes, we demonstrated that HMGB1, acting in the nucleus, exerts a potent repressive effect on LXRα activity and hepatic lipogenesis during metabolic stresses, such as F/R or HFD feeding, suggesting a protective role on the development of NAFLD.

The nuclear role of HMGB1 might be more complex than initially envisioned and may depend on cell type, nature of environmental signals, and the pathophysiological context. In the context of metabolic stress, we demonstrate *in vitro*, using primary culture of hepatocyte, that HMGB1 exerts its repressive effect on lipid metabolism in a cell-autonomous manner, thus supporting a model where HMGB1 remains inside the hepatocyte. One can presume that either HMGB1 stays in the nucleus and/or translocates in the cytoplasm. Our ChIP-seq data clearly showed that upon the nutritional challenges we have applied, HMGB1 leaves the chromatin, exemplified by reduced binding affinity to DNA and loss of TSS occupancy, triggering a number of changes in gene transcription. Other studies have described a similar impairment of DNA affinity by HMGB1 in cells subjected to stress (*26*, *40*). In a recent study, it was shown that in senescent cells, HMGB1 leaves the nucleus leading to a significant change in gene expression (mostly up-regulation) and in chromatin topology (*26*), which is in agreement with our results in hepatocytes. Despite being poorly documented, it has also been described that HMGB1 in the nucleus may both be bound and unbound to DNA, and that even when unbound it may still reside within the nucleus during cell cycle (*40*). This supports a model where upon stressors or outside signals, HMGB1 may dissociate from DNA but stays in the nucleus. Yet, the precise mechanisms regulating this biological event and the role of unbound HMGB1 within the nucleus remain unknown, and further experiments are required to understand the underlying mechanism. At the same time, the channeling of HMGB1 between nucleus and cytoplasm is determined by a variety of post-translational modifications such as acetylation, methylation or phosphorylation. During inflammatory challenges for example, acetylation has been described to regulate the accessibility of the HMGB1 nuclear localization signal to the cargo proteins, thus balancing the protein pool between nucleus and cytoplasm (*37*). In the context of a metabolic stress, it has been suggested that the histone deacetylase SIRT1, a key metabolic sensor (*41*), may play a significant role in the acetylation status of HMGB1 and its sub-cellular localization (*42*).

Our data suggest that in response to micro-environmental signals, HMGB1 may dissociate from the chromatin thus affecting biological functions, including metabolic processes. On CD, we found HMGB1 occupying 134 gene loci belonging to metabolic functions, which have been identified as depending on the activity of LXRα. As LXRα is a key lipogenic transcription factor involved in cholesterol metabolism and liver lipogenesis, the de-repression of its activity induced by HMGB1 deletion logically translates into liver steatosis (*31*, *32*). The molecular mechanism behind the inhibition of the hepatic lipogenesis by HMGB1 is still not entirely clear. The immediate mechanism and the simplest scenario would be a direct or indirect binding of HMGB1 with LXRα, even though a direct physical interaction was not seen in our co-immunoprecipitation assay (**Fig. 7B**). One cannot rule out that using more sensitive techniques, a physical interaction might be found as a physical interactions of HMGB1 with transcription factors have been described, notably sterol regulatory element-binding proteins (SREBPs) and the glucocorticoid receptor (GR) (*24*, *25*). Study of the HMGB1-interactome in hepatocytes *in vivo* might be interesting to explore, albeit technically challenging.

Our ATAC-seq data helped to demonstrate that chromatin compaction was not regulated by HMGB1 under CD and during the nutritional challenges (fig. S7), suggesting that the HMGB1-mediated repression was likely not mediated through a nucleosomal re-organization. This hypothesis was important to test, as several reports demonstrated a key role of HMGB1 in the nucleosome arrangement remodeling associated to transcription modulation *in vitro* (*22*). At least in the *in vivo* context of liver steatosis, our results support a minor role for HMGB1 in regulating nucleosomal landscapes, which represents a significant layer of epigenetic control of transcription. However, our ChIP-seq data suggested DNA occupancy as a likely mechanism of repression. HMGB1 has a very high level of DNA occupation in the basal state and that it is located equally in the promoter region, CDS and distal intergenic region. However, upon metabolic stress, HMGB1 appears to leave the chromatin, particularly the TSS regions (**Fig. 7C**). This suggests that HMGB1 DNA occupancy is correlated with changes in gene transcription, but interestingly, the occupancy rate in the TSS is not necessarily related to the level of repression, as shown by two equally-repressed genes (*Acly* and *Fasn*) with heterogeneous TSS occupation (fig. S9B-C). Hence, occupancy appears to be an important factor, but likely not the only one. Of note, our data using inducible *Hmgb1* deletion via AAV8-TBG-Cre show that the absence of HMGB1 consistently leads to the up-regulation of genes involved in hepatic lipogenesis, suggesting a causal relationship between HMGB1 and gene expression (**Fig. 7E-F**). These results are corroborated by a study of Sofiadis et al, depicting a map of HMGB1 binding genome-wide in senescent cells using a combination of RNA-seq, ChIP-seq and Hi-C (chromatin conformation capture). Interestingly, in primary cells at senescent state, HMGB1 leaves the chromatin, triggering profound changes in chromatin dynamics and gene transcription, in a similar fashion as seen by us. Additionally, Hi-C data demonstrated that HMGB1 binds to TAD (Topology Associated Domain) boundaries, known to regulate chromatin topology and consequently gene expression. In addition to this paper, a recent study has also evoked an RNA-binding property as a another functional layer for HMGB1 to regulate gene expression (*26*, *43*). Therefore, 3-D conformation and RNA binding clearly represent additional mechanisms by which HMGB1 could mediate its repressive effect on LXRα, which is therefore worthwhile to further investigate in the context of liver steatosis.

Overall our study helped to uncover HMGB1-mediated LXRα repression as new mechanism modulating liver lipogenesis during metabolic stress. Boosting these functions of HMGB1 may constitute a new therapeutic approach to counteract the deleterious effect of enhanced LXRα activity in patients with NAFLD.

## Materials and Methods

### Experimental Design

This study aimed to decipher the precise role of the nuclear factor HMGB1 in hepatocytes during metabolic stress. For this, a cell specific knockout mice model where *Hmgb1* gene is deleted specifically in hepatocytes (HMGB1^ΔHep^) and its control counterpart (HMGB1^fl/fl^) were subjected to nutritional stress such as high fat diet and fasting/refeeding. A combination of OMICS studies has been employed to nail down the potential mechanism behind HMGB1 repressive effect on hepatic lipogenesis such as microarray, ATAC-seq or ChIP-seq. All studies identified lipid metabolism as a key function and transcription factor LXRα as a key piece that might be repressed by HMGB1. *In vivo* studies using adenovirus-mediated shRNA expression targeting LXRα were employed to functionally test the interdependence of HMGB1 and LXRα. *In vitro* assays were used to measure how HMGB1 could regulate the transcriptional activation using specific responsive elements (RE)-containing luciferase reporter. For in vivo studies, adult age-matched Cre +/− carrying *Hmgb*1 floxed gene called HMGB1^ΔHep^ mice and their control Cre −/− carrying *Hmgb*1 floxed gene named HMGB1^fl/fl^ littermates were co-housed to reduce variability. Animal numbers for each study type were determined by the investigators on the basis of data from previous similar experiments or from pilot studies. For OMICS studies, displayed animals were chosen as representative from the whole cohort: (i) for the microarray 4 animals per genotype/per challenge, (ii) 2 animals per genotype/per challenge for the Chip-seq and (ii) 2 animals per genotype/per challenge for the ATAC-seq have been analyzed. For neutral lipid analysis and histology experiments, sample identities were not known in most cases and were randomized. For *in vitro* studies, at least three biological replicates were used in three separate experiments.

### Mouse Phenotyping

Breeding and experimental procedures were performed in accordance with institutional guidelines for animal research and were approved by the Animal Care and Use Ethics Committee US006 CREFRE - CEEA-122 (protocol 17/1048/03/20). Animals were housed in temperature and humidity controlled facilities under a 12 hour-light period with free access to food and water. All animals were aged between 2 to 3 months at the beginning of the experimentations. Hepatocyte-specific deletion of *Hmgb1* gene noted HMGB1^ΔHep^ were generated crossing Alb-CRE^+/−^ (Jackson Laboratory, Ban Harbor, ME, USA) with *Hmgb1* floxed mice noted HMGB1^fl/fl^ (a generous gift from Dr. Robert F. Schwabe, Columbia University, NY, USA), littermates Alb-CRE^−/−^ HMGB1^Flox/Flox^ (HMGB1^fl/fl^) were used as control. Hepatocyte-specific deletion of *Pparγ* gene noted PPARγ ^ΔHep^ were generated crossing Alb-CRE^+/−^ (Jackson Laboratory, Ban Harbor, ME, USA) with *Pparγ* floxed mice noted PPARγ^fl/fl^ (a generous gift from Pr. W.A Wahli, University of Lausanne, Switzerland), littermates Alb-CRE^−/−^ PPARγ^Flox/Flox^ (PPARγ^fl/fl^) were used as control. At the time of sacrifice, tissues and organs were dissected, weighted and directly snap frozen in liquid nitrogen and stored at −80°C.

### Genotyping

DNA extraction and PCR were performed using Kapa mouse genotyping kit (Kapa Biosystems, Wilmington, MA, USA) according to the manufacturer protocol. PCR reactions were performed using following primers: Alb-CRE: 5′-ACCGGTCGATCGAAACGAGTGATGAG-3 (forward) and 5′-AGTGCGTTCGAACGCTAGAGC-3′ (reverse), LoxP1 5′-TAAGAGCTGGGTAAACTTTAGGTG-3′ (forward) and 5′-GAAACAGACAAGCTTCAAACTGCT-3′ (reverse), LoxP2 5′-TGACAGGATACCCAGTGTTAGGGG-3′ (forward) and 5′-CCAGAGTTTAATCCACAGAAGAAA-3′ (reverse).

### Interventional experiments

– For diet induced-obesity experiments, mice were fed with a normal chow diet (CD, Research Diets, New Brunswick, NJ, USA) or a high fat diet (HFD60%, Research Diets, New Brunswick, NJ, USA) for 12 or 24 weeks. To induce liver steatosis, mice were subjected to HFD60% with 30% fructose (Sigma-Aldrich, Saint Louis, MO, USA), dissolved in the drinking water or choline deficient diet supplemented with 60% fat (CD-HFD60%, Research Diets, New Brunswick, NJ, USA). For the fasting-refeeding, mice under normal chow diet (CD) were starved 6 hours from Zeitgeber 14 (ZT14) and refeed for 8 hours with the CD and 20% glucose (Sigma-Aldrich, Saint Louis, MO, USA) in the drinking water.
– Body composition was assessed using the EchoMRI (Echo Medical Systems, Houston, TX, USA).
– Indirect calorimetry was performed after 24 h of acclimatization in individual cages. Oxygen consumption, carbon dioxide production, and food and water intake were measured (Phenomaster; TSE Systems, Bad Homburg v.d.H, Germany) in individual mice at 15-min intervals during a 24-h period at constant temperature (22°C). The respiratory exchange ratio ([RER] = *V*co_2_/*V*o_2_) was measured. The glucose oxidation (in g/min/kg^0.75^ = [(4.545 × *V*co_2_) − (3.205 × *V*o_2_)]/1000) and lipid oxidation (in g/min/kg^0.75^ = [1.672 × (*V*o_2_ − *V*co_2_)]/1000) were calculated. Ambulatory activities of the mice were monitored by infrared photocell beam interruption (Sedacom; Panlab-Bioseb).
– For *Hmgb1* gene deletion at adult age, HMGB1^fl/fl^ male mice at 8 weeks of age were injected intravenously (i.v) with 10^11^ genomic copies per mouse with adeno-associated-virus (AAV8) containing a liver-specific promoter, thyroxine-binding globulin (TBG) promoter driving either GFP or Cre recombinase (Penn Vector Core, University of Pennsylvania, PA, USA) to generate control mice noted AAV-GFP or liver specific HMGB1 knockout noted AAV-CRE. 7 days after injections, animals were euthanized.
– To knockdown LXR, adult male HMGB1^fl/fl^ and HMGB1^ΔHep^ mice (8-12 week-old) were injected i.v with an adenovirus expressing an shRNA targeting LXRα (kindly provided by Dr Catherine Postic, Cochin Institute, Paris, France). For both adenovirus protocols, 10^13^ adenoviral infectious particles were diluted in 0.9% NaCl and administered in a total volume of 100 μl per animal. 7-10 days after injection, control (scramble RNA noted sh*SCR*) and sh*LXRα* expressing mice were subjected to fasting/refeeding challenges as described previously. To study HFD-induced liver steatosis, mice were first subjected to a 4 week-HFD60%, then injected with sh*SCR* and sh*LXRα*, and mice were euthanized 7-10 post injections.
– For Insulin acute injection, CD or HFD60% fed HMGB1^fl/fl^ and HMGB1^ΔHep^ mice were fasted for 16 hours and then injected i.p (intra-peritoneal) with 0,75U/kg of human insulin and mice were sacrificed 15 minutes later.
– For LXR *in vivo* activation, synthetic agonist T0901317 (30mg/kg, Bertin Bioreagent, Montigny le Bretonneux, France) was administered orally by four consecutive daily gavages on 8-week-old HMGB1^fl/fl^ and HMGB1^ΔHep^ adult male mice. Mice were starved one hour before the fourth gavage, and maintained starved for 5 more hours before euthanasia.
– For hepatic VLDL-triacylglycerol production assay, 8-week-old HMGB1^fl/fl^ and HMGB1^ΔHep^ adult male mice fasted overnight received an intravenous injection of 10% tyloxapol (500 mg/kg) (Sigma-Aldrich, T8761, Saint Louis, MO, USA). Blood was collected from the tail vein at 0, 1, 2, 3 and 4 hours for triglyceride assays.

### Glucose/Insulin/pyruvate tolerance test

– Glucose (GTT), Insulin (ITT) and pyruvate (PTT) tolerance tests were performed under chow diet or after 12 weeks of HFD after an overnight fast. Glucose (Sigma, G8270, Saint Louis, MO, USA) was orally administered at 1.5 g/kg dose, Insulin was injected i.p at 0.75 U/kg and pyruvate (Sigma, P2256, Saint Louis, MO, USA) was administrated by i.p. injection at 1.5g/kg. For all tolerance tests the glycaemia evolution was then monitored at the tail vein using Accu-Check glucometer (Roche). Plasma insulin (Mercodia, Upasal, Sweden) was determined by ELISA in the fasted state or at indicated times.

### Primary Hepatocyte Isolation

Mouse hepatocytes were isolated as previously described via 2-step collagenase perfusion as described by Fortier *et al* (*44*). Hepatocytes were allowed to attach for 90 minutes on collagen-coated plates in RPMI containing 10% FBS (Gibco), followed by overnight starvation in serum-free medium before experiments (Lipogenesis and β-oxidation assay).

### Lipogenesis assays

– For *in vitro* measurement, one day after isolation, primary hepatocytes were serum-starved for 3 hours and incubated for 3-hour with [1-^14^C] acetate (1 μCi/ml; Perkin Elmer, Boston, MA) and 5.5 mM of non-labeled (cold) glucose in DMEM medium. At the end of incubation, cells were washed twice with cold PBS 1X and harvested into 0.25 ml of 0.1% SDS for subsequent protein measurement and total lipid extraction with 1 ml of chloroform/methanol (2v/1v). Lipid extracts were washed with 70% ethanol, and then dissolved into chloroform/methanol (2v/1v). Radioactivity was measured on a multipurpose scintillation counter (LS 6500; Beckman Coulter). All assays were performed in duplicates, and data normalized to cell protein content.
– For *in vivo* measurement of lipogenesis activity, animals were fasted for 6 hours at ZT14 and received an i.p. bolus of 2 mg/g glucose containing 0.4μCi/g of [3-3H]-D-glucose (Perkin-Elmer, NET331C, Waltham, MA, USA). After 1 hour, liver, epididymal, subcutaneous and brown adipose tissues were collected and snap-frozen in liquid nitrogen.
– For palmitate oxidation assay: Cells were preincubated for 3 hours with ^14^Cpalmitate (1uCi/mL; Perkin Elmer, Boston MA) and non labeled (cold) palmitate. Palmitate was coupled to a fatty acid-free BSA in a molar ratio of 5:1. Following incubation, ^14^CO_2_ and ^14^C-ASM were measured as previously described (*45*). Briefly, assayed medium was transferred into a custom-made Teflon 48-well trapping plate. The plate was clamped and sealed, and perchloric acid was injected through the perforations in the lid into the medium, which drives CO_2_ through the tunnel into an adjacent well, where it was trapped in 1N NaOH. Following trapping, the media was spun twice and ^14^C-ASM measured by scintillation counting. Aliquots of NaOH and medium were transferred into scintillation vials, and radioactivity was measured on a multipurpose scintillation counter (LS 6500; Beckham Coulter). All assays were performed in triplicates, and data were normalized to protein content.

### Liver neutral lipid analysis

Hepatic lipids were extracted by the “Folch” procedure before being quantified using mass spectrometry. Briefly, 50mg of liver were homogenized in 1mL water:methanol (1:2 v/v), 5 mM EGTA. Lipids are then extracted using a methanol: chloroform: water (2.5:2.5 : 1.7 v/v) mix. After a solid phase extraction, purification and desiccation, all lipids are eluted in ethyl-acetate and analyzed by a gas chromatography combined with mass spectrometry (GC-MS) (ISQ Thermo).

### Microarray Gene Expression Studies

Gene expression profiles were performed at the GeT-TRiX facility (GénoToul, Génopole Toulouse Midi-Pyrénées) using Agilent Sureprint G3 Mouse GE v2 microarrays (8×60K, design 074809) following the manufacturer’s instructions. For each sample, Cyanine-3 (Cy3) labeled cRNA was prepared from 200 ng of total RNA using the One-Color Quick Amp Labeling kit (Agilent) according to the manufacturer’s instructions, followed by Agencourt RNAClean XP (Agencourt Bioscience Corporation, Beverly, Massachusetts). Dye incorporation and cRNA yield were checked using Dropsense™ 96 UV/VIS droplet reader (Trinean, Belgium). 600 ng of Cy3-labelled cRNA were hybridized on the microarray slides following the manufacturer’s instructions. Immediately after washing, the slides were scanned on Agilent G2505C Microarray Scanner using Agilent Scan Control A.8.5.1 software and fluorescence signal extracted using Agilent Feature Extraction software v10.10.1.1 with default parameters.

### Microarray data statistical analysis

Microarray data were analyzed using R (*46*) and Bioconductor packages (*47*). Raw data (median signal intensity) were filtered, log2 transformed and normalized using the quantile method (*48*) with the limma package (*49*).

A model was fit using the limma lmFit function (*49*). Pairwise comparisons between biological conditions were applied using specific contrasts. In cases where Agilent has multiple probe sequences for the same gene, the probe with the best p-value was selected. Probes with a p-value ≤ 0.01 were considered to be differentially expressed between conditions.

Normalized log intensities were averaged (n == 4) within each group and heatmaps were generated with the ComplexHeatmap package (*50*). Venn diagrams were generated with the Vennerable package (https://github.com/js229/Vennerable). Functional pathway enrichment was performed in R using the hypergea package’s hypergeometric test (https://cran.r-project.org/package=hypergea). GO annotations were obtained using biomaRt (*51*) and the graphite package (*52*) was used to obtain pathways from the Reactome database. ChEA (https://doi.org/10.1093/bioinformatics/btq466) was interrogated via the Enrichr website (*53*) and tabular results were imported into R. Barcharts were constructed using ggplot2 (*54*). The network of pathways largely shared between F/R and HFD was constructed in R as csv files that were imported into Cytoscape (*55*).

### ChIP-seq

Briefly, frozen liver biopsies (100-200 mg) harvested from HMGB1^fl/fl^ and HMGB1^ΔHep^ mice under CD, upon HFD60% or after F/R, were minced and fixed at room temperature in PBS-1% formaldehyde (Sigma-Aldrich, 47608, Saint Louis, MO, USA) for 20 minutes. After sonication, chromatin immunoprecipitation was performed using anti-HMGB1 antibody (Abcam, ab18256, Cambridge, UK). Immunoprecipitated DNA was subjected to library preparation and single-end sequencing on a NextSeq 500 at EMBL GeneCore (Heidelberg, Germany).

### ATAC-seq

Flash-frozen liver biopsies were sent to Active Motif to perform the ATAC-seq assay. The tissue was manually dissociated, isolated nuclei were quantified using a hemocytometer, and 100,000 nuclei were tagmented as previously described (*56*), with some modifications based on (*57*) using the enzyme and buffer provided in the Nextera Library Prep Kit (Illumina). Tagmented DNA was then purified using the MinElute PCR purification kit (Qiagen), amplified with 10 cycles of PCR, and purified using Agencourt AMPure SPRI beads (Beckman Coulter). Resulting material was quantified using the KAPA Library Quantification Kit for Illumina platforms (KAPA Biosystems), and sequenced with PE42 sequencing on the NextSeq 500 sequencer (Illumina).

### ATAC-seq and ChIP-seq data analysis

ATAC-seq and ChIP-seq reads were first mapped to the mouse genome UCSC build hmm10 using Bowtie2 2.2.8 (*58*). Aligned reads were then filtered to keep only matched pairs and uniquely mapped reads. Peaks were called with MACS2 2.2.1 (*59*) algorithm using a mappable genome size of 2.73e^9^. To process ChIP-seq datasets, MACS2 was run with the “Delta” genotype as a negative control as in this condition the HMGB1 protein expression is reduced by 90 % and signal detected in “Delta” libraries, defined as background noise, was substracted from the “Flox” libraries. ATAC-seq datasets were processed without a control file and with the -nomodel option. Called peaks that were on the ENCODE blacklist of known false ChIP-seq peaks were removed. Signal maps and peak locations were used as input to the statistical analysis performed with the R package ChIPseeker (*60*). DESeq2 (*61*) was used to identify differential binding sites and differential open chromatin profiles. Motifs and GO enrichment analysis were respectively performed using JASPAR (*62*) and the R package ReactomePA (*63*).

### Histology

Tissue samples were fixed in 10% formalin (Sigma-Aldrich, HT501128, Saint Louis, MO, USA) for 24 hours, then incubated at 4°C in 70% ethanol before being paraffin-embedded or in 30% sucrose before being cryo-embedded with Tissue-Tek OCT (Sakura FineTek Europe, Alphen aan den Rijn, The Netherlands). Paraffin embedded livers were sliced at 5 μm. For Periodic Acid Schiff reaction, sections are incubated in 0.5% periodic acid in water for 5 minutes then transferred to Schiff reagent (Sigma-Aldrich, 3952016, Saint Louis, MO, USA) for 15 minutes. Sections were counterstained with Mayer’s hematoxylin (Sigma-Aldrich, MHS16, Saint Louis, MO, USA) before mounting. Liver-cryo sections were post-fixed with 10% formalin 15 minutes prior staining with Oil-red-O (Sigma-Aldrich, MHS16, Saint Louis, MO, USA)(60% solution in isopropanol-Sigma-Aldrich, 33539, Saint Louis, MO, USA). After counter-staining with hematoxylin, slides are mounted with aqueous mounting media. Stained slides were scanned using a Nanozoomer scanner (Hamamatsu Photonics, Hamamatsu City, Japan). Images quantification was performed using Image J freeware (NIH, USA).

### Western blotting

Tissues were homogenized in RIPA buffer (TRIS 20 mM, NaCl 150 mM, EDTA 1 mM, EGTA 1 mM, TRITON X100 1%, Tetra-Sodium Pyrophosphate 2.5 mM, B-Glycerophosphate 1 mM, Sodium orthovanadate 1 mM) containing protease and phosphatase inhibitors (Sigma-Aldrich, St. Louis, MO, USA) using Precellys sample lyzer (Bertin Technologies, Montigny le Bretonneux, France). Western blots were performed using standard procedures using antibodies against HMGB1 (1:1000, ab18256, Abcam, Cambridge, UK), Phospho-AKT S473 (1:1000, CST 4060, Cell Signaling Technology, Danvers, MA, USA), total AKT (1:1000, CST 9272, Cell Signaling Technology, Danvers, MA, USA), HA (1:1000, CST 3724 Cell Signaling Technology, Danvers, MA, USA), Myc-tag (1:1000, CST 2276, Cell Signaling Technology, Danvers, MA, USA) and GAPDH (1: 2000, ab181602, Abcam, Cambridge, UK), was used as a loading control.

### Reporter assay

For reporter assay, Ad293 cells were cultured in 96 well plates with DMEM containing 10% FB Essence (Avantor Seradigm, USA) and transfected using Transit-LT1 (Mirus Bio, Madison, WI, USA) with plasmid encoding 4 LXR response elements fused with luciferase, human HA-LXR (HA-hLXR) and RXR. HMGB1 plasmid was purchased from Origene. 24h after transfection, cells media was changed to DMEM containing 2% charcoal striped and dialyzed media with 0.1 uM of T0901317 and/or 1uM of LG100268 (noted LG268) (Cayman Chemical, USA). After overnight treatment, luciferase activity was assayed using a luciferase assay system (Promega, USA). Bioluminescence was quantified using a luminometer and normalized to β-Gal activity.

### Co-immunoprecipitation

Ad293 cells were plated in 6 well-plate and transfected as previously described with 1 ug of HA-hLXR and/or HMGB1 plasmids. 24h after transfection, cells were treated with 0.1uM of T0901317 overnight. Cells were lysed in IP buffer (20mM Tris-HCl pH8, 100mM NaCl, 0.1%NP40, 10% glycerol, 2uM PMSF and 1mM DTT) supplemented with antiprotease and antiphosphatase cocktails.

IP was performed using HA-conjugated beads (Sigma) for 2h at 4°C, following wash step, beads were resuspended in 2X Laemmli buffer and western blot was performed as previously described.

### Gene expression

RNA were extracted using GenJET RNA purification kit (ThermoScientific, Waltham, MA, USA) and DNAse treatment (Qiagen, Hilden, Germany). After dosage with Xpose (Trinean, Gentbrugge, Belgium) reverse transcription was performed using High Capacity cDNA reverse transcription kit (Applied Biosystem, Foster City, CA, USA) according to the manufacturer protocol. Real-Time -qPCR was performed with indicated primer pairs gene expression is normalized using *36b4* reference gene expression. Primer sequences are available in **Table S3**.

### Microfluidic qPCR

Expression analyses of lipogenesis related-genes **(Table S3**) were performed by quantitative PCR with Fluidigm Biomark^®^ technology (Genome & Transcriptome GenoToul Platform). First-strand cDNA templates were pre-amplified with Preamp Master Mix (Fluidigm) and reactions were achieved in a Fluidigm Biomark® BMK-M-96.96 plate according to the manufacturer’s recommendations. Relative gene expression values were determined using the 2^−ΔΔCT^ method. The expression analyses data are an average of seven individuals for HMGB1^fl/fl^ mice and 10 individuals for HMGB1^ΔHep^ mice. As described before, the *36B4* gene expression levels were used for data standardization.

### Plasma analysis

Whole blood is drawn out from the inferior vena cava after euthansia, and plasma is prepared after centrifugation (5 minutes; 4 °C; 8000 rpm). Circulating AST (ASpartate aminoTransferase) and ALT (ALanine aminoTransferase) levels were determined in plasma by the Phénotypage-CREFRE facility using a Pentra400 biochemical analyzer (HORIBA Medical, Kyoto, Japan). HMGB1 circulating levels were assessed by ELISA (ST51011, IBL International, Hamburg, Germany) on 10 uL of plasma, according to the manufacturer guidelines.

### Statistics

Analyses are performed using GraphPad Prism 7 (GraphPad Software, La Jolla, CA, USA). Potential outliers were identified using ROUT algorithm (GraphPad Software) and removed from analysis. All data are expressed as mean ± SEM, except otherwise indicated. Statistical significance was determined by Mann & Withney, one-way ANOVA or two-way ANOVA followed by a Tuckey post-hoc test. P values <0.05 were considered significant (*p < 0.05; **p < 0.01; ***p < 0.001; ****p<0,0001).

## H2: Supplementary Materials

Fig. S1. Metabolic explorations of Hepatocyte specific *Hmgb1* deleted mice subjected to chow-diet.

Fig. S2. Metabolic explorations of Hepatocyte specific *Hmgb1* deleted mice subjected to high-fat diet.

Fig. S3. Hepatocyte specific *Hmgb1* deleted mice exhibit a severe liver steatosis upon various diets.

Fig. S4. Hierarchical clustering and color heatmap of differentially expressed gene comparing livers of HMGB1^fl/fl^ and HMGB1^ΔHep^ mice.

Fig. S5. Hepatocyte specific *Pparγ* deletion does not modify liver steatosis in mice after F/R challenge.

Fig. S6. Hepatocyte specific *Hmgb1* deleted mice exhibit a severe liver steatosis under metabolic stressors that is restored by knocking-down LXRα *in vivo*.

Fig. S7. Hepatocyte specific *Hmgb1* deletion does not remodel chromatin.

Fig. S8. Validation of HMGB1 ChIP *in vitro* and *in vivo.*

Fig. S9. Genomic and around TSS (−1kb/+1kb) region distribution of HMGB1.

Fig. S10. Successful *in vivo* knockdown of *Hmgb1* gene and protein using AAV-TBG-Cre.

**Table S1. List of genes highly occupied by HMGB1 in Chow Diet compare to HFD**

**Table S2. List of genes highly occupied by HMGB1 in Chow Diet compare to F/R**

**Table S3. Primers for Real time qPCR**

## Acknowledgments

The authors would like to thank the phenotyping facility (Anexplo-US006/CREFRE, Toulouse, France) for all plasma analysis and technical assistance. We also thank the Histology (Lucie Fontaine) and the Functional Biochemistry (Alexandre Lucas) Facilities at the i2MC for technical assistance (UMR1048-Toulouse, France). We thank Claire Naylies and Yannick Lippi for their contribution to microarray fingerprints acquisition and microarray data analysis carried out at GeT Genopole Toulouse Midi-Pyrénées facility (https://doi.org/10.15454/1.5572370921303193E12). Jean-José Maoret (GeT-Santé, UMR1048-Toulouse, France) for his technical help during the microfluidic PCR experiments, Justine Bertrand-Michel and the lipidomic facility (Genotoul-MetaToul Lipidomique, UMR1048-Toulouse, France) for all lipid analysis and technical assistance. We kindly thank Dr. Pierre-Damien Denechaud (i2MC-INSERM-Toulouse) for fruitful discussions and kind advices.

## Funding

This study is supported by grants from INSERM, Paul Sabatier University, the Agence Nationale de la Recherche (ANR-17-CE14-0016, J-PP) and Association Française d’Etude et de Recherche sur l’Obésité (J-PP). JP is supported by a scholarship from Paul Sabatier University, EP is supported by a scholarship from Agence Nationale de la Recherche (ANR-17-CE14-0016) and RP is supported by a scholarship from Région Midi-Pyrénées-INSERM (n°15050341).

## Author contributions

JP designed research, performed all experiments, analyzed data and wrote the manuscript; AD, AP, AB, SD, EP, RP and EM performed experiments and analyzed data, J.SI and JM analyzed high-throughput data and draft related figures. TC and GL carried out the ChIP-seq experiments, analyzed data and reviewed the manuscript. AM and W.AW generously provided PPAR*γ*^fl/fl^ and PPAR*γ*^ΔHep^ mice. W.AW reviewed and commented on the manuscript. R.FS kindly provided HMGB1 floxed mice and reviewed the manuscript. AY and IC-L have provided valuable inputs, daily support, have reviewed and edited the manuscript. CP and FB kindly provided the adenovirus expressing shRNA-targeting LXR, reviewed and commented on the manuscript; CM performed lipogenesis and Beta-Ox assays *in vitro* and *in vivo*, analyzed data and reviewed and commented on the manuscript; HG designed and performed experiments, analyzed data provided reagents, gave important input related to the study design, reviewed and commented on the manuscript, PV and CD have provided daily support, fruitful discussions, fundings and revised and commented on the manuscript; J-PP conceived the original hypothesis, designed all experiments, performed experiments, analyzed data, wrote the manuscript, provided fundings and supervised the project.

## Competing interests

The authors declare no conflicts of interest.

## Data and materials availability

All data needed to evaluate the conclusions in the paper are present in the paper and/or the Supplementary Materials.

## Notes

### Competing Interest Statement

The authors have declared no competing interest.

