## Supplementary Figures for "Nuclear HMGB1 protects from non-alcoholic fatty liver diseases through negative regulation of liver X receptor": Supplementary Materials Personnaz J et al .pdf

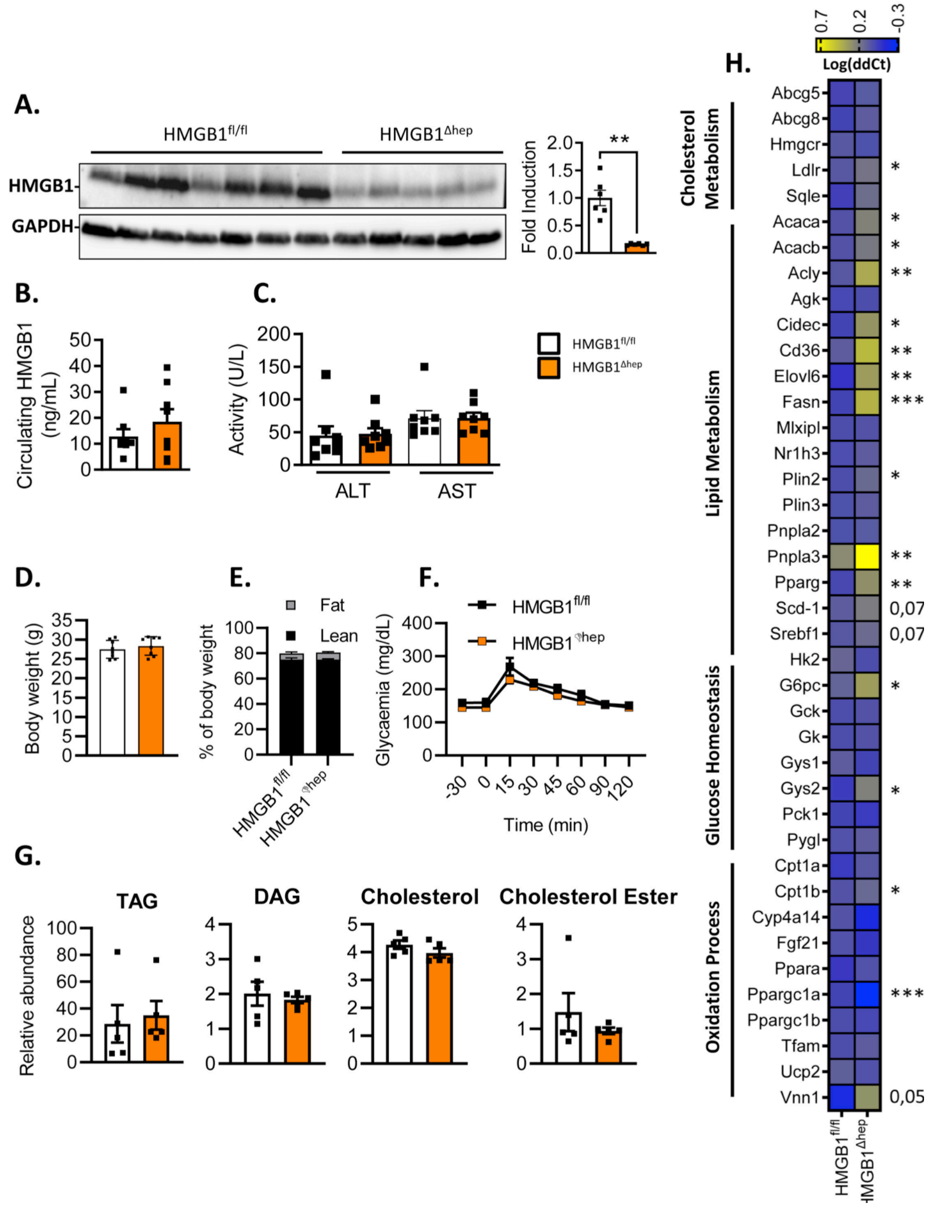


**Fig. S1. Metabolic explorations of Hepatocyte specific *Hmgb1* deleted mice subjected to chow-diet.**

(**A**) Representative immunoblot targeting HMGB1. GAPDH was used as a loading control. Densitometric quantification (right panel) performed on the whole animal cohort, on liver biopsies from on 8 week-old HMGB1$^{fl/fl}$ (n=5) and HMGB1$^{\Delta Hep}$ (n=5) mice fed on chow diet. (**B-C**) Circulating levels of HMGB1 (**B**) and hepatic transaminases (**C**), (**D**) Body weight, (**E**) percentage of lean and fat mass, (**F**) analysis of oral glucose tolerance test and (**G**) liver neutral lipid levels in liver sections of 8 week-old HMGB1$^{fl/fl}$ and HMGB1$^{\Delta Hep}$ mice fed on chow diet. (**H**) Heatmap showing gene expression of specific markers involved in hepatic lipid and glucose metabolism determined by microfluidic RT-qPCR. Data are means ± SEM of three independent experiments. *p<0.05, **p<0.01, ***p<0.001, ****p<0.0001 by unpaired Mann and Whitney comparison

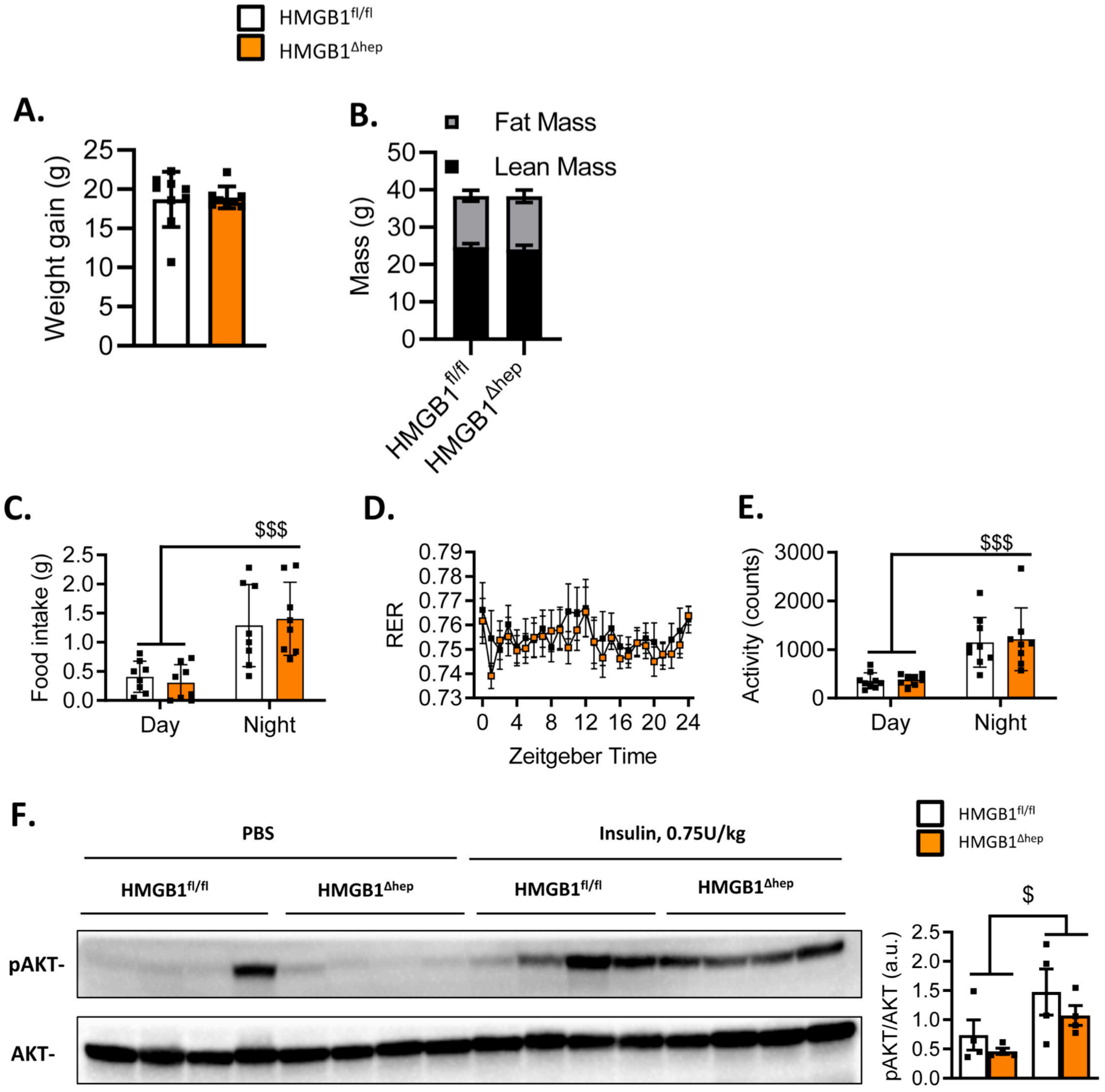


**Fig. S2. Metabolic explorations of Hepatocyte specific *Hmgb1* deleted mice subjected to high-fat diet.**

(**A**) Weight gain and (**B**) body fat and lean mass of HMGB1$^{fl/fl}$ (n=9) and HMGB1$^{\Delta Hep}$ (n=8) mice on HFD. (**C-E**) Indirect calorimetry analysis monitoring (**C**) food intake, (**D**) Respiratory exchange Ratio (RER) and (**E**) spontaneous physical activity of HMGB1$^{fl/fl}$ (n=8) and HMGB1$^{\Delta Hep}$ (n=8) mice on HFD during 24 hours. (**F)** Representative immunoblot targeting p-AKT and Tot-AKT was used as a loading control. Densitometric quantification (right panel) performed on the whole animal cohort, on liver biopsies from on 8 week-old HMGB1$^{fl/fl}$ and HMGB1$^{\Delta Hep}$ mice fed on chow diet and challenged with an acute injection of PBS (100uL) (n=4 for each genotype) or insulin (i.p, 0.75 U/kg) (n=4 for each genotype). Livers were harvested 15 minutes after insulin stimulation. Data are means ± SEM of three independent experiments. *p<0.05, **p<0.01, ***p<0.001, ****p<0.0001 by unpaired Mann and Whitney comparison or two-way ANOVA

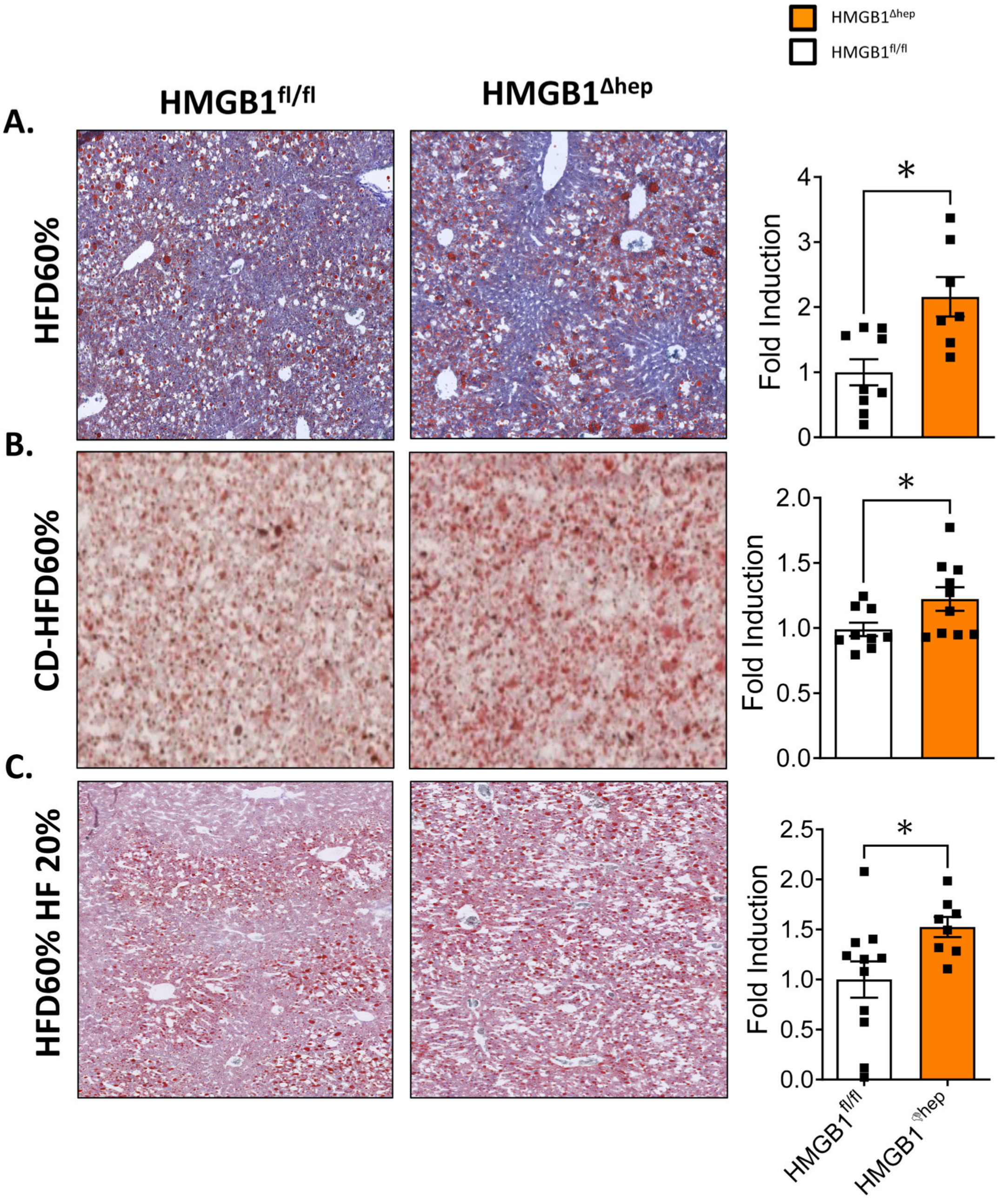


**Fig. S3. Hepatocyte specific *Hmgb1* deleted mice exhibit a severe liver steatosis upon various diets.**

Adult $HMGB1^{fl/fl}$ and $HMGB1^{\Delta Hep}$ mice were subjected to various numbers of diets and liver steatosis was assessed by Oil Red-O staining on liver section: (**A**) 24 week-HFD60% (respectively n=9 and n=7), (**B**) 4 week-CD-HFD60% (respectively n=9 and n=10), and (**C**) 12 week-HFD60%-high fructose (HF) (respectively n=11 and n=8). Data are means ± SEM of three independent experiments. *p<0.05, **p<0.01, ***p<0.001, ****p<0.0001 by unpaired Mann and Whitney comparison or two-way ANOVA.

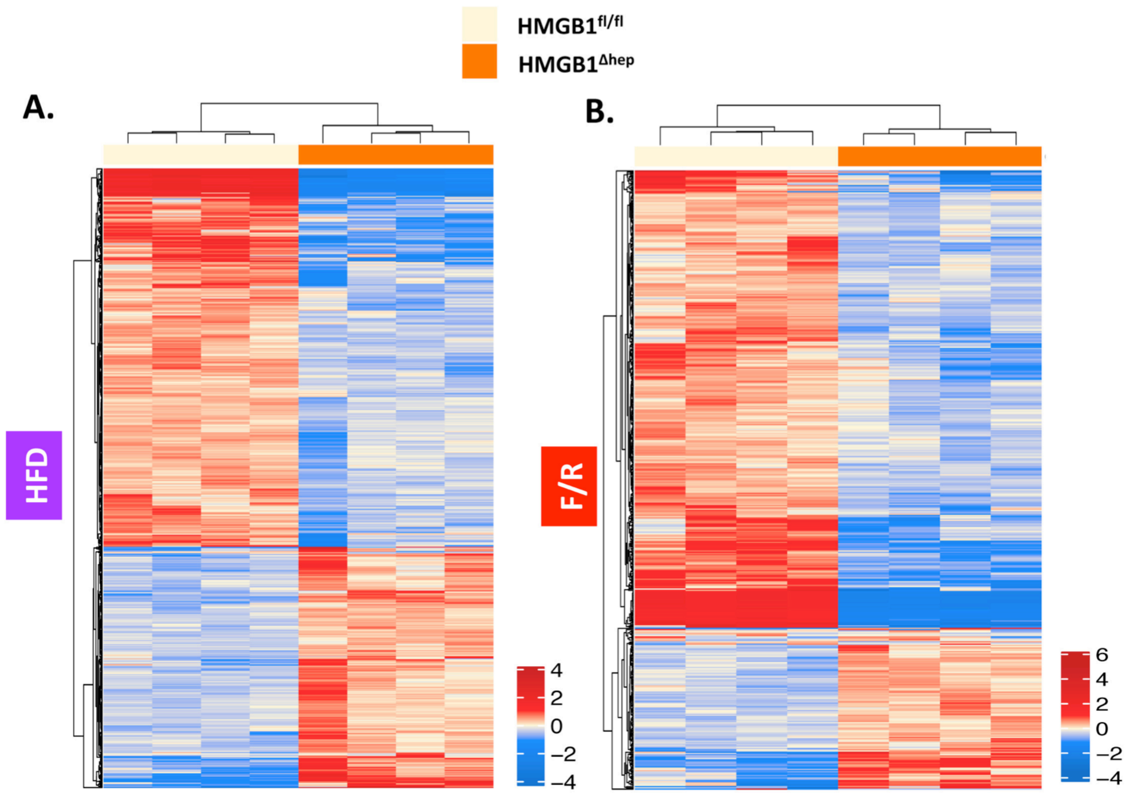


**Fig. S4. Hierarchical clustering and color heatmap of differentially expressed genes between livers of HMGB1$^{fl/fl}$ and HMGB1$^{\Delta Hep}$ mice.**

(**A-B**) Heatmap displaying genes that are differentially expressed in the livers of HMGB1$^{\Delta Hep}$ mice compared to HMGB1$^{fl/fl}$ mice (fold change > 1.5; P-Value <= 0.01) from (**A**) HFD and (**B**) F/R regimes, where each track represents one animal (n=4 per genotype). Values are normalized expression, centered by row means.

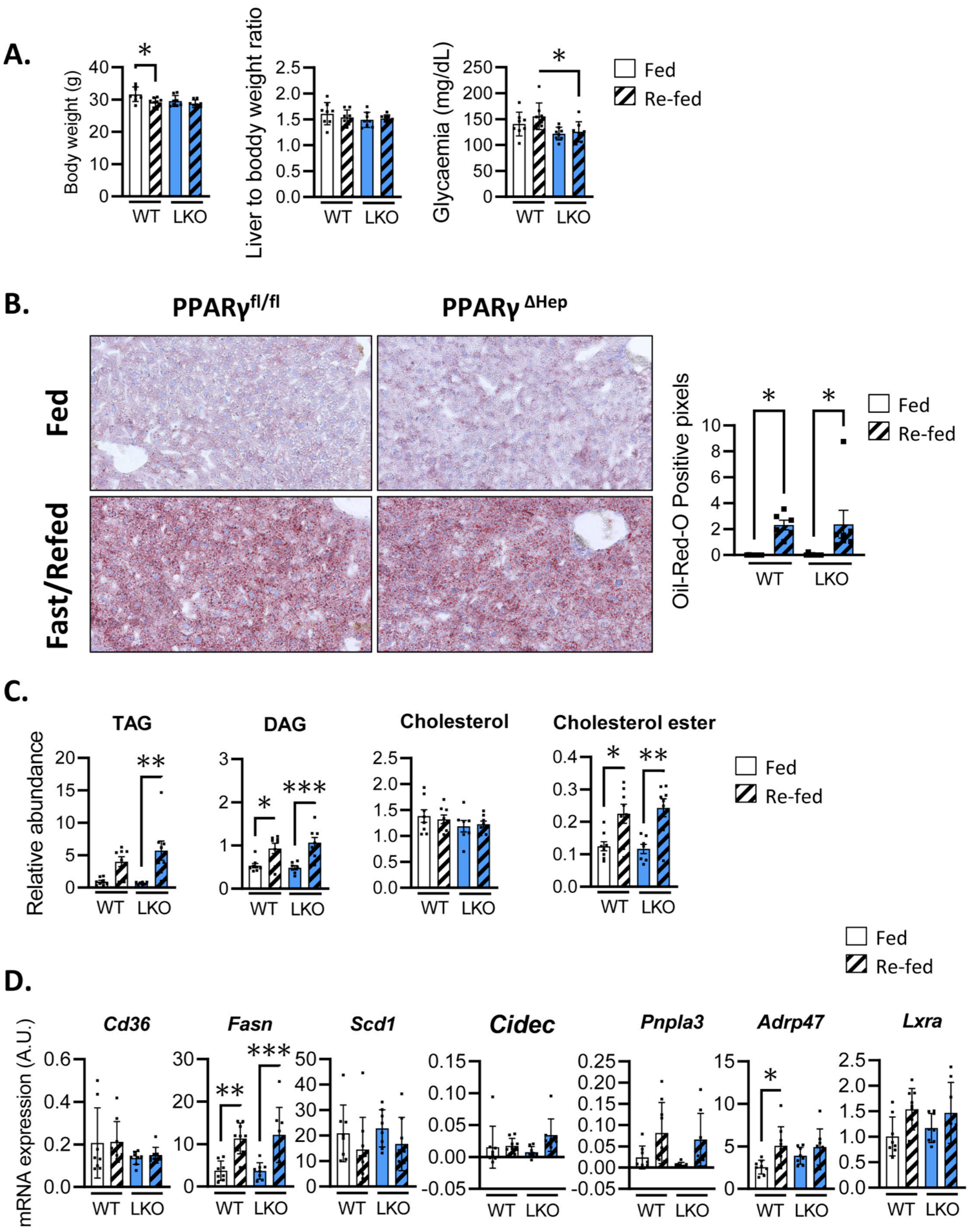


**Fig. S5. Hepatocyte specific *Pparγ* deletion does not modify liver steatosis in mice after F/R challenge.**

PPARγ $^{fl/fl}$ and PPARγ $^{\Delta Hep}$ (n=8) mice were subjected to F/R challenge. (**A**) Body weight, liver/body weight ratio and starved glycaemia of mice on fed (6 hours)-refeed (8 hours). Hepatic steatosis was assessed by (**B**) Oil-Red O staining with quantification (right panel) and (**C**) neutral lipids analysis. (**D**) mRNA expression of hepatic steatosis markers from liver biopsies was determined by RT-qPCR. Data are means ± SEM of three independent experiments. *$p<0.05$, **$p<0,01$, ***$p<0.001$, ****$p<0.0001$ by unpaired Mann and Whitney comparison.

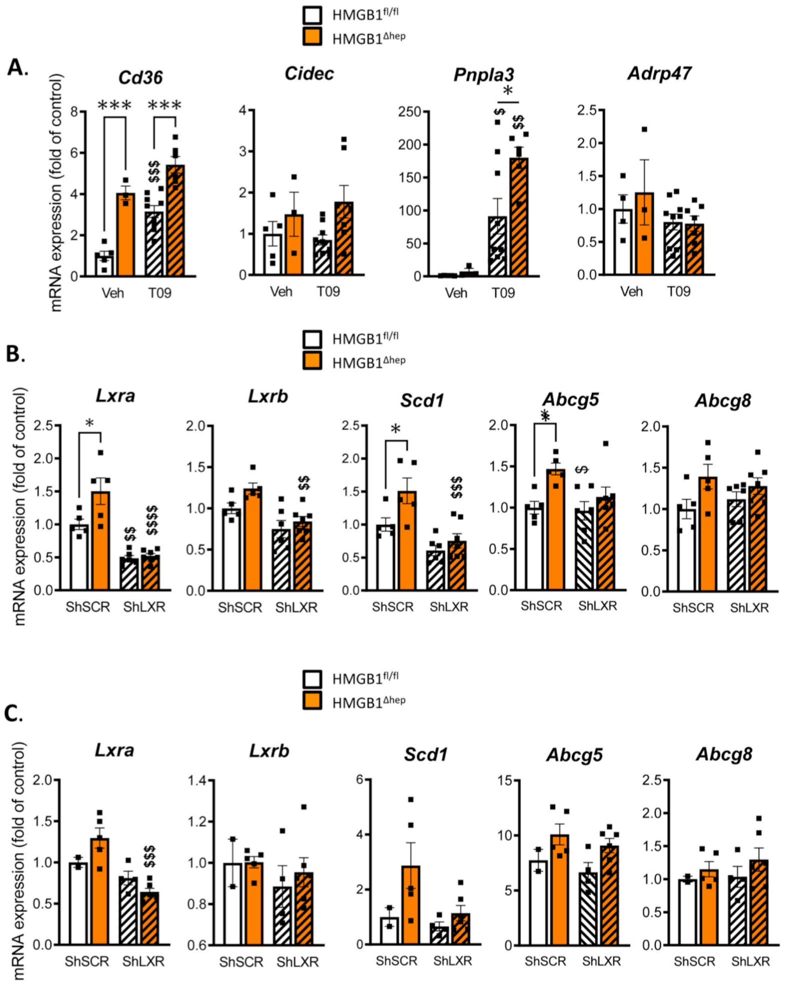


**Fig. S6. Hepatocyte specific *Hmgb1* deleted mice exhibit a severe liver steatosis under metabolic stress that is restored by knocking-down LXRα *in vivo*.**

(**A**) 8 week-old HMGB1$^{fl/fl}$ (n=5) and HMGB1$^{\Delta Hep}$ (n=9) mice fed on chow diet were treated either with vehicle (5% carboxy-methyl-cellulose) (HMGB1$^{fl/fl}$ (n=5) and HMGB1$^{\Delta Hep}$ (n=3)) or LXRα synthetic agonist T0901317 (oral gavage, 30 mg/kg/day) (HMGB1$^{fl/fl}$ (n=9) and HMGB1$^{\Delta Hep}$ (n=6)) for four consecutive days, after 6 hours starvation on the last day mice were sacrificed. (**A**) mRNA expression of hepatic steatosis markers from liver biopsies of HMGB1$^{fl/fl}$ and HMGB1$^{\Delta Hep}$ mice after a fasting/refeeding challenge. (**B**) HMGB1$^{fl/fl}$ and HMGB1$^{\Delta Hep}$ mice were infected with either adenovirus expressing a scramble (shSCR, HMGB1$^{fl/fl}$, n=6 and HMGB1$^{\Delta Hep}$, n=7) or an LXRα shRNA (shLXR, HMGB1$^{fl/fl}$, n=5 and HMGB1$^{\Delta Hep}$, n=5) sequence, then subjected 7 days later to a F/R challenge was then subjected to RT-qPCR analysis of the indicated LXRα dependent genes. (**C**) HMGB1$^{fl/fl}$ and HMGB1$^{\Delta Hep}$ mice were subjected to HFD for four weeks and then infected with either adenovirus expressing a scramble (shSCR, HMGB1$^{fl/fl}$, n=2 and HMGB1$^{\Delta Hep}$, n=5) or an *Lxra* shRNA (sh*Lxra*, HMGB1$^{fl/fl}$, n=4 and HMGB1$^{\Delta Hep}$, n=6) sequence and euthanized 7 days later. (**B-C**) Liver steatosis and sh*Lxra* efficiency was assessed by measuring indicated LXRα dependent gene expression using RT-qPCR. Data are means ± SEM. *p<0.05, **p<0.01, ***p<0.001, ****p<0.0001 for HMGB1$^{fl/fl}$ and HMGB1$^{\Delta Hep}$ comparison, by unpaired Mann and Whitney comparison. $ p<0.05, $$ p<0.01, $$$ p<0.001, for treatment effect by one-way ANOVA.

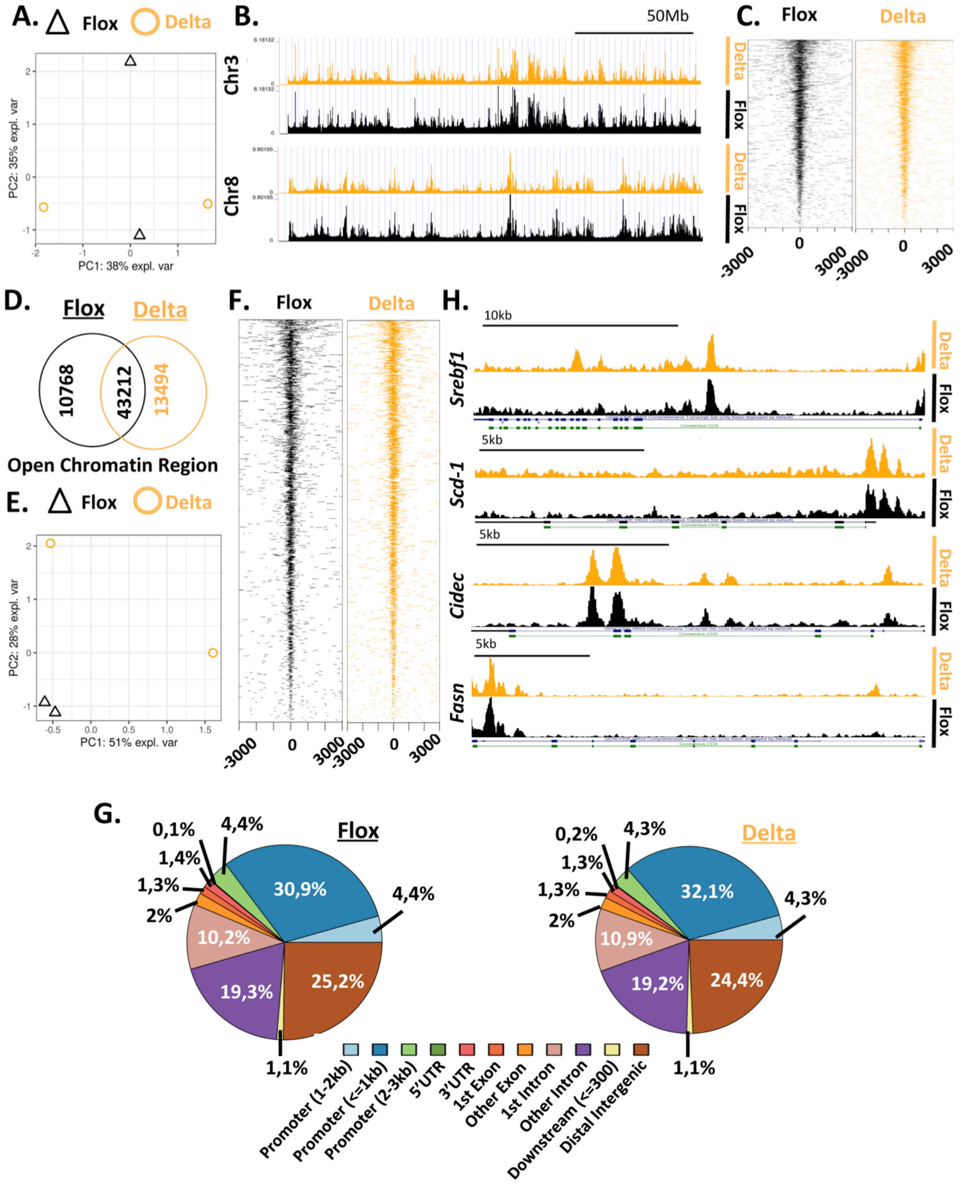


**Fig. S7. Hepatocyte specific *Hmgb1* deletion does not remodel chromatin.**
(**A**) Principal component analysis scores plot of ATAC-seq data of liver tissue from HMGB1$^{fl/fl}$ (black triangle, n=2) and HMGB1$^{\Delta Hep}$ (orange circle, n=2)) mice on chow diet. (**B**) UCSC genome browser of tracks showing HMGB1 similar chromatin open region in Chromosome 3 and 8. (**C**) Average coverage around TSS in liver biopsies of HMGB1$^{fl/fl}$ (black) and HMGB1$^{\Delta Hep}$ (orange) mice. (**D**) Venn Diagram showing overlapping opened region and and (**E**) Principal component analysis scores plot of ATAC-seq data of liver tissue from HMGB1$^{fl/fl}$ (black triangle, n=2)) and HMGB1$^{\Delta Hep}$ (orange circle, n=2) mice subjected to F/R. (**F**) Average coverage around TSS in liver biopsies of HMGB1$^{fl/fl}$ (black) and HMGB1$^{\Delta Hep}$ (orange) mice after F/R challenge. (**G**) Chart pie displaying the genomic distribution of open chromatin domain in liver biopsies from HMGB1$^{fl/fl}$ (black, n=2)) and HMGB1$^{\Delta Hep}$ (orange, n=2) mice. (**H**) UCSC genome browser snapshots of tracks showing HMGB1 chromatin open region of canonical LXRα responsive gene loci (*Srebf1*, *Scd-1*, *Cidec*, *Fasn*) in liver biopsies of HMGB1$^{fl/fl}$ (black) and HMGB1$^{\Delta Hep}$ (orange) mice after F/R challenge.

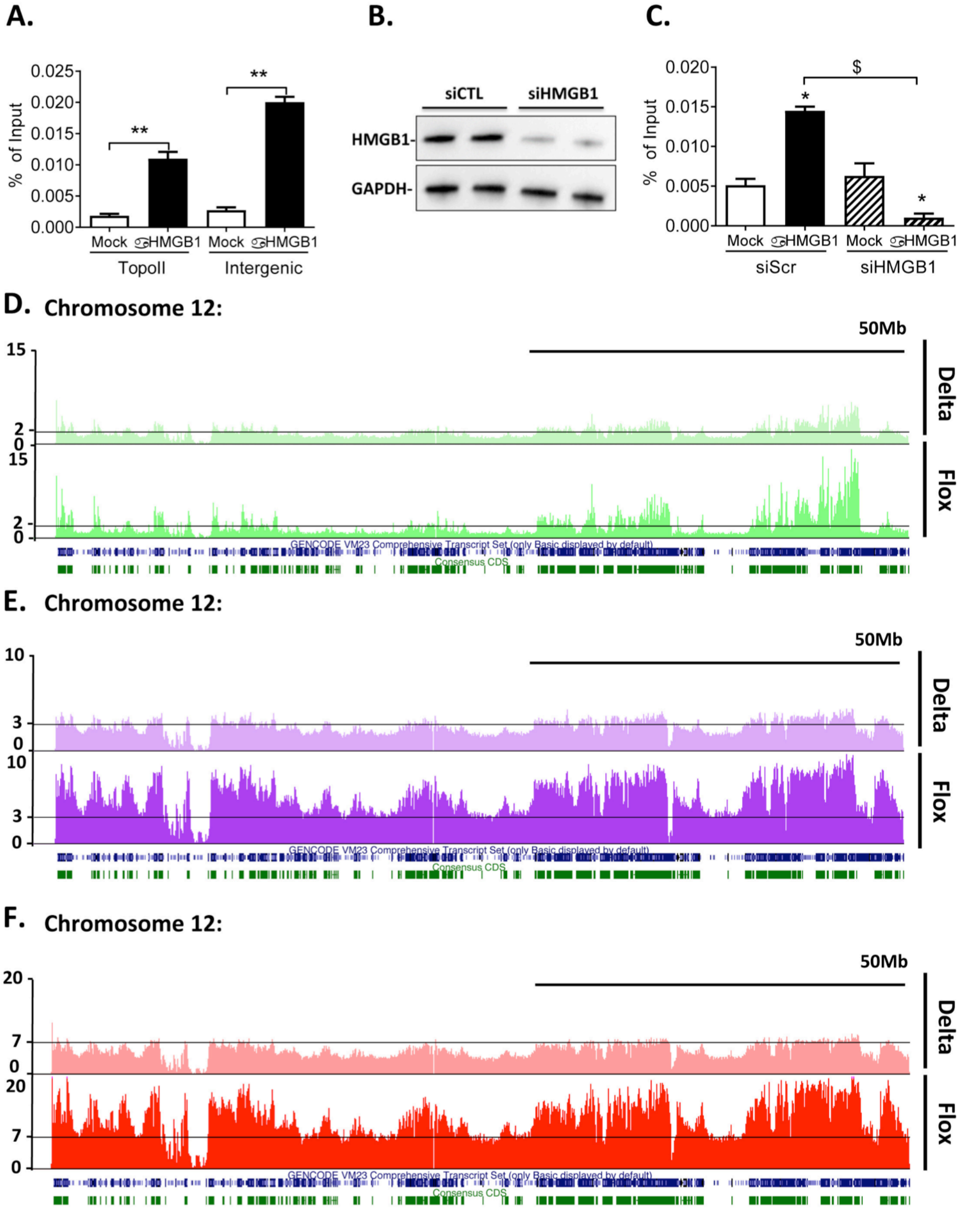


**Fig. S8. Validation of HMGB1 ChIP *in vitro* and *in vivo*.**
(**A**) Chromatin immuno-precipitation was performed against HMGB1 on U2OS cells and enrichment on TOPO isomerase II promoter sequence as well as intergenic region were measured by RT-qPCR. (**B-C**) To validate specificity of the ChIP, (**B**) U2OS cells were first transfected with a siRNA targeting HMGB1 and after successful knockdown of HMGB1 showed by immunoblot targeting HMGB1 with GAPDH as a loading control, (**C**) ChIP experiment has been conducted, and siHMGB1 treated cells showed no DNA enrichment of TOPOII promoter region while siScramble exhibits a significant DNA enrichment. (**D-F**) UCSC genome browser of tracks from the ChIP-seq data, comparing the sequencing signal in Chromosome 2 from HMGB1$^{fl/fl}$ (n=2) and HMGB1$^{\Delta Hep}$ (n=2) mice upon (**D**) Chow diet (green), (**E**) HFD (purple) and (**F**) F/R (red). Data are means ± SEM of three independent experiments (**A-C**) *p<0.05 ** p<0.01 αHMGB1 vs Mock, **$**<0.05 siScramble vs siHMGB1 by unpaired Mann and Whitney comparison or two-way ANOVA

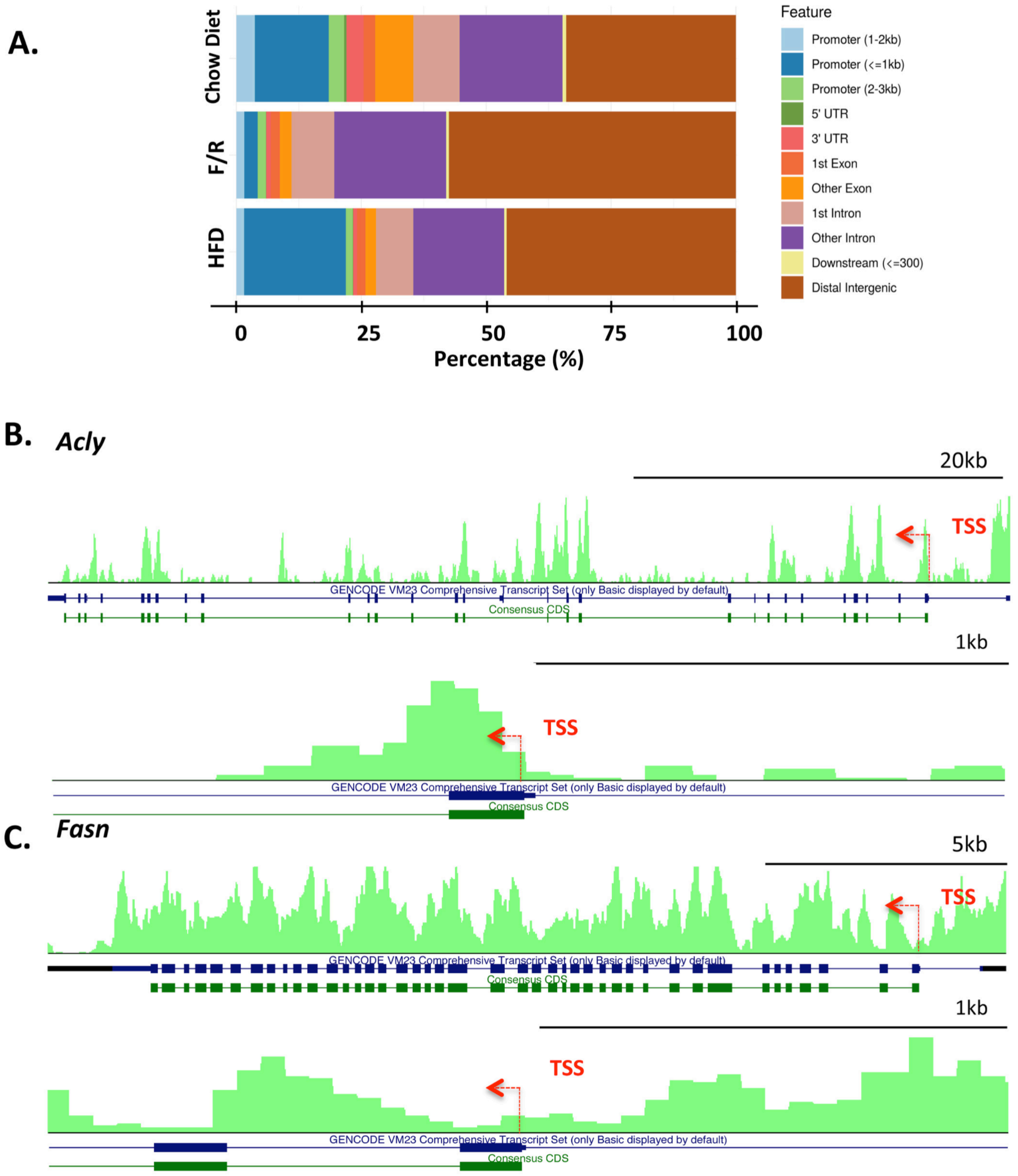


**Fig. S9. Genomic and around TSS (-1kb/+1kb) region distribution of HMGB1.**
(**A**) Genomic distributions of enriched regions identified in ChIP-seq data sets in liver biopsies of HMGB1$^{fl/fl}$ mice subjected to chow diet, HFD and F/R. (**B-C**) HMGB1 coverage on gene locus and TSS region (-1kb/+1kb) of LXRα responsive gene: *Acly* (**B**) and *Fasn* (**C**).

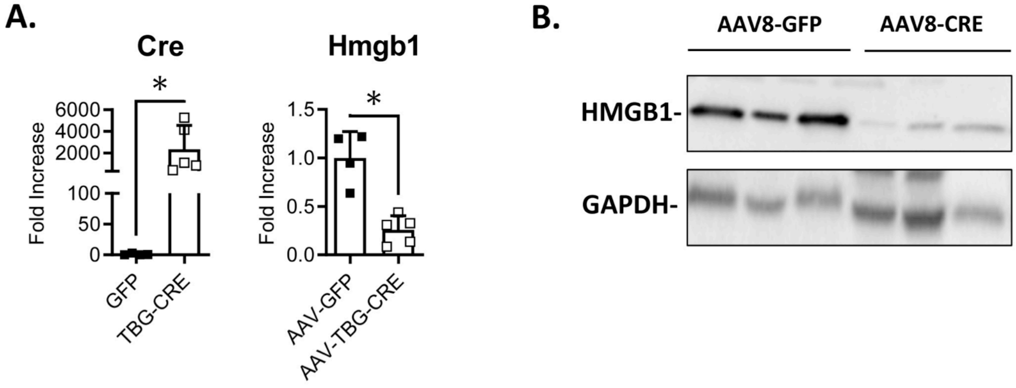


**Fig. S10. Successful *in vivo* knockdown of *Hmgb1* gene and protein using AAV-TBG-Cre.** Adult $HMGB1^{fl/fl}$ mice were infected either with AAV8-Gfp (n=3) or AAV8-Tbg-Cre (n=3) to selectively generate *Hmgb1* deletion in hepatocytes *in vivo.* (**A**) The level of expression of *Hmgb1* gene was determined using RT-qPCR in whole liver. (**B**) Representative immunoblot targeting HMGB1 in liver extracts. GAPDH was used as a loading control. Data are means ± SEM. *p<0.05, **p<0.01, ***p<0.001, ****p<0.0001 by unpaired Mann and Whitney comparison.

**Table S1. List of genes highly occupied by HMGB1 in Chow Diet compare to HFD**

| GO « Integration of Energy Metabolism » | GO « Phospholipid Metabolism » |
|---|---|
| Abcc8 | Acp6 |
| Acaca | Agpat1 |
| Acacb | Agpat5 |
| Acly | Arf1 |
| Acsl3 | Awat2 |
| Adcy5 | Cds2 |
| Adipor2 | Cept1 |
| Adra2c | Cpne1 |
| Cacna1a | Cpne3 |
| Cacna1c | Cpne7 |
| Cacna2d2 | Ddhd2 |
| Cacnb2 | Enpp6 |
| CD36 | Etnk1 |
| Fasn | Gnpat |
| Gcg | Gpat2 |
| Gcgr | Gpat3 |
| Glp1r | Gpat4 |
| Gna11 | Hadha |
| Gna14 | Inpp4b |
| Gna15 | Inpp5f |
| Gnai1 | Liph |
| Gnai2 | Lpcat2 |
| Gnaq | Lpcat3 |
| Gnas | Lpin3 |
| Gnb1 | Mfsd2a |
| Gnb2 | Mgll |
| Gnb3 | Miga1 |
| Gng11 | Mtm1 |
| Gng12 | Mtmr14 |
| Gng13 | Mtmr7 |
| Gng3 | Osbpl10 |
| Gng5 | Osbpl5 |
| Gng8 | Osbpl8 |
| Itpr1 | Phospho1 |
| Itpr2 | Pik3ca |
| Itpr3 | Pik3cd |
| Kcnb1 | Pik3r2 |
| Kcnc2 | Pik3r3 |
| Kcng2 | Pip5k1a |
| Kcnj11 | Pip5k1b |
| Marcks | Pitpnb |
| Mlx | Pitpnm2 |
| Mlxipl | Pla2g12a |
| Plcb1 | Pla2g15 |
| Plcb2 | Pla2g1b |
| Plcb3 | Pla2g2d |
| Prkag2 | Pla2g2e |
| Prkar1a | Pla2g4d |
| Prkar1b | Pla2g4f |
| Prkca | Pla2g5 |
| Rapgef4 | Plb1 |
| Slc2a1 | Plekha1 |

| | |
|---|---|
| | Plekha2 |
| | Plekha5 |
| | Pnpla6 |
| | Pnpla8 |
| | Ptpn13 |
| | Rab4a |
| | Selenoi |
| | Slc44a1 |
| | Slc44a2 |
| | Slc44a3 |
| | Slc44a5 |
| | Stard10 |
| | Stard7 |
| | Taz |
| | Tnfaip8l1 |
| | Tnfaip8l2 |
| | Tnfaip8l3 |
| | Tpte |
| | Vac14 |

### Table S2. List of genes highly occupied by HMGB1 in Chow Diet compare to F/R

| GO « Integration of Energy Metabolism » | GO « Phospholipid Metabolism » |
|---|---|
| Acaca | Acp6 |
| Acacb | Agpat1 |
| Acly | Agpat5 |
| Acsl3 | Arf1 |
| Acsl4 | Awat2 |
| Adcy5 | Cds2 |
| Adcy6 | Cept1 |
| Adipor2 | Cpne1 |
| Adra2c | Cpne3 |
| Agpat1 | Cpne7 |
| Cacna1a | Ddhd2 |
| Cacna1c | Enpp6 |
| Cacna2d2 | Etnk1 |
| Cacnb2 | Gnpat |
| Cacnb3 | Gpat2 |
| Cd36 | Gpat3 |
| Fasn | Gpat4 |
| Gcg | Hadha |
| Gcgr | Inpp4b |
| Glp1r | Inpp5f |
| Gna11 | Liph |
| Gna14 | Lpcat2 |
| Gna15 | Lpcat3 |
| Gnai1 | Lpin3 |
| Gnai2 | Mfsd2a |
| Gnaq | Mgll |
| Gnas | Miga1 |
| Gnb1 | Mtm1 |
| Gnb2 | Mtmr14 |
| Gnb3 | Mtmr7 |
| Gng10 | Osbpl10 |
| Gng11 | Osbpl5 |
| Gng12 | Osbpl8 |
| Gng13 | Phospho1 |
| Gng3 | Pik3ca |
| Gng5 | Pik3cd |
| Itpr1 | Pik3r2 |
| Itpr2 | Pik3r3 |
| Itpr3 | Pip5k1a |
| Kcnb1 | Pip5k1b |
| Kcnc2 | Pitpnb |
| Kcng2 | Pitpnm2 |
| Kcns3 | Pla2g12a |
| Marcks | Pla2g15 |
| Mlx | Pla2g1b |
| Mlxipl | Pla2g2d |
| Plcb1 | Pla2g2e |
| Plcb2 | Pla2g4d |
| Plcb3 | Pla2g4e |
| Prkab2 | Pla2g4f |
| Prkaca | Pla2g5 |
| Prkag2 | Plb1 |

| | |
|---|---|
| Prkar1a | Plekha1 |
| Prkar1b | Plekha2 |
| Prkar2b | Plekha5 |
| Prkca | Pnpla6 |
| Rap1a | Pnpla8 |
| Rapgef3 | Ptpn13 |
| Rapgef4 | Rab4a |
| Slc2a1 | Selenoi |
| | Slc44a1 |
| | Slc44a2 |
| | Slc44a3 |
| | Slc44a5 |
| | Stard10 |
| | Stard7 |
| | Taz |
| | Tnfaip8l1 |
| | Tnfaip8l2 |
| | Tnfaip8l3 |
| | Tpte |
| | Vac14 |

### Table S3. Primers for Real time qPCR

| Gene | Forward Primer | Reverse Primer |
|---|---|---|
| *36B4* | AGTCGGAGGAATCAGATGACGAT | GGCTGACTTGGTTGCTTTGG |
| *Abcg5* | TCGCAACGGTCATTTTCA | GCCAAAAGAGCAGCAGAGAAATA |
| *Abcg8* | GCGTCTGTCGATGCTGGTC | ATCCATTGGCCACCCTTGT |
| *Acc1* | CTCCTTTGCCTTCCGACATC | TACCATGCCAATCTCATTTCCTC |
| *Acc2* | TGGAGGCAACAGGGTCATAGA | CGCAGCGATGCCATTGT |
| *Acly* | AAAGCTTGGCCTCGTCGG | GGGACGAAGGGTTCAATGAGA |
| *Adrp47* | GTGTCGTCGTAGCCGATGC | CCATTTCTCAGCTCCACTCCAC |
| *Agk* | GTGTTTGGCAACCAGCTCATT | GCTGCGGGATTGAGAAAGAC |
| *Fabp4* | TTCGATGAAATCACCGCAGA | GGTCGACTTTCCATCCCACTT |
| *Cd11b* | TCGGACGAGTTCCGGATTC | TGTGATCTTGGGCTAGGGTTTC |
| *Cd11c* | GATTTCAGCATCCCAGATCCC | CCAGATCCACCAGTCCATCC |
| *Cd36* | GGACATACTTAGATGTGGAACCCATA | TGTTGACCTGCAGTCGTTTTG |
| *Cd45* | ACATGCTGCCAATGGTTCTG | GTCCCACATGACTCCTTTCCTATG |
| *Chrebp* | ACATCAGCGCTTTGACCAGAT | TGCGCGTCCGGACATAG |
| *Cidec* | GACTTTATTGGCTGCCTGAACG | ATCTCCTTCACGATGCGCTT |
| *Cpt1a* | CACCAACGGGCTCATCTTCT | CCTTCTATCGAATTTGCTCTGGTT |
| *Cpt1b* | GTGCAAGCAGCCCGTCTAG | TTGCGGCGATACATGATCA |
| *Cre* | CATTGCTGTCACTTGGTCGT | CATTTGGGCCAGCTAAACAT |
| *Cyp4a14* | TCAGTCTATTTCTGGTGCTGTTC | GAGCTCGTTGTCCTTCAGATGGT |
| *Elovl6* | TCTGATGAACAAGCGAGCCA | TGGTCATCAGAATGTACAGCATGT |
| *F4/80* | TGACAACCAGACGGCTTGTG | GCAGGCGAGGAAAAGATAGTGT |
| *Fasn* | ATCCTGGAACGAGAACACGATCT | AGAGACGTGTCACTCCTGGACTT |
| *Fgf21* | GGTCAAGTCCGGCAGAGGTA | CTGTTCCATCCTCCCTGATCTC |
| *G6pc* | ACGTATGGATTCCGGTGTTTG | CAGCTGCACAGCCCAGAA |
| *Gck* | TCGCAGGTGGAGAGCGA | TCGCAGTCGGCGACAGA |
| *Gk* | GTGTTTGGCAACCAGCTCATT | GCTGCGGGATTGAGAAAGAC |
| *Gys1* | CGGCTTTGGCTGCTTTATG | CCAGAATGTAAATGCCGTAAGCT |
| *Gys2* | CCAACGACGGATGGCTTTAA | GATCCTGACGGAGAAGGTGGTA |
| *Hk2* | CGCCGGATTGGAACAGAA | CCCGTCGCTAACTTCACTCACT |
| *Hmgb1* | AGCCCTGTCCTGGTGGTATT | CCAGGCAAGGTTAGTGGCTA |
| *Hmgcr* | TTGTTCACGCTCATAGTCGCT | GACACATCTTCGTCCAGACCC |
| *L-pk* | AGTCGTGCAATGTTCATCCCT | TCGACTCAGAGCCTGTGGC |
| *Ldl-r* | TGGATCCACCGCAACATCTA | CTCTTTACGCCCTTGGTGTCA |
| *Lpl* | TTATCCCAATGGAGGCACTTTC | CACGTCTCCGAGTCCTCTCTCT |
| *Lxra* | AGGAGTGTCGACTTCGCAAA | CTCTTCTTGCCGCTTCAGTTT |
| *Lxrbeta* | AAGCAGGTGCCAGGGTTCT | TGCATTCTGTCTCGTGGTTGT |
| *Pck1* | ATGTTCGGGCGGATTGAAG | TCAGGTTCAAGGCGTTTTCC |
| *Pnpla2* | CGCCTCTCGAAGGCTCTCTT | TGTAGCCCTGTTTGCACATCTC |
| *Pnpla3* | AGCCCGTCTCTGATGCACTT | ACGCGGTCACCTTCGTGT |
| *Ppara* | TGGCAGCAATATCAGAGGTAGATTC | ATCATATCAAAGGAGCTGCCAAA |
| *Pparg* | CCGAAGAACCATCCGATTGA | TTTGTGGATCCGGCAGTTAAG |
| *Pparg1ca* | AAAGGATGCGCTCTCGTTCA | GGAATATGGTGATCGGGAACA |
| *Ppargc1b* | TTGAGGTGTTCGGTGAGATTGTAG | GAAGGTGATAAAACCGTGCTTCTG |
| *Pygl* | CAAGTGTCCCAAGAGGGTGT | TGTAATGTTCGGCCCATGTA |
| *Scd-1* | CAACACCATGGCGTTCCA | GGTGGGCGCGGTGAT |

| | | |
|---|---|---|
| *Serpin2* | CAACACAGGGATCCAGGTCT | CATGAGGCCGTGACTTGATG |
| *Sqle* | CTGCACTTGGTTGGTTTCTGAC | GGAGGCTACCGTGTTCTCCA |
| *Srebf1* | CTGGCTTGGTGATGCTATGTTG | GACCATCAAGGCCCCTCAA |
| *Tfam* | GCACCCTGCAGAGTGTTCAA | CGCCCAGGCCTCTACCTT |
| *Tip47* | GGCTGGACAGACTGCAGGA | TCTTGAGCCCCAGACACTGTAG |
| *Ucp2* | CCTCAAAGCAGCCTCCAGAA | TCAATCGGCAAGACGAGACA |
| *Vnn1* | ATGAGGTTTATGCCTTTGGAGC | CCACAGGTGCGTAAATTGGTAG |
